# Elevated polyadenylated histone gene transcription in human cancer

**DOI:** 10.64898/2026.10.14.758961

**Authors:** Ronald M. Paranal, Terri D. Bryson, Guillermina Rios Rios, Kevin K. Nguyen, Kami Ahmad, Steven Henikoff

**Affiliations:** Basic Sciences Division, Fred Hutchinson Cancer Center, Seattle, WA 98109, United States; Division of Medical Oncology, Department of Medicine, University of Washington, Seattle, WA 98195, United States; HHMI, Chevy Chase, MD 20815, United States

## Abstract

Histone production is tightly restricted to S-phase to prevent genomic instability, yet human cancers with an overabundance of RNA Polymerase II (RNAPII) at histone genes are associated with aggressive cancers (1). To better understand histone gene overexpression in cancer, we applied a quantitative PCR assay to measure total replication-coupled histone output across the histone gene clusters using cell line models. We observed pervasive accumulation of polyadenylated histone transcripts, supported by analysis of >1,400 RNA-sequencing datasets in human cancers, with more than 70% of patients showing tumor-specific increases. This alternative processing preferentially enriches for histone H2B, which lacks replication-independent variants, across >8,300 tumors comprising diverse cancer types and transcriptional subtypes. We also found that disruption of nuclear components required to maintain RNAPII levels at the histone genes increased polyadenylated histone transcripts. Our results suggest that S-phase-dependent histone genes are subject to unscheduled gap-phase transcription in human cancer.

## Main

Genome-wide hypertranscription in cancer portends a poor prognosis (2, 3) and fuels histone overexpression that may contribute to abnormal proliferation and whole-arm chromosomal loss associated with cancer recurrence (1, 4). Transcription of the multicopy histone genes that reside in the major and minor histone clusters is coordinated during S-phase to synthesize canonical replication-coupled histone mRNA (5–7). However, whether RNAPII generates transcripts beyond the constraints of this highly regulated process in cancer remains unclear, as previous studies have not measured the total output of the human replication-coupled histone genes. Here we show the accumulation of replication-coupled histone transcripts across human cancers, with histone H2B disproportionally enriched. Thus, cancer-associated hypertranscription during gap phase may broadly support cancer initiation and proliferation through elevated histone synthesis, while uniquely fulfilling the required replacement for histone H2B.

To measure total transcriptional output of all replication-coupled histone genes, we optimized degenerate PCR primers. The amino acid sequences of these genes are largely invariant and highly conserved, yet coding sequences of these histone genes are considerably variable (8, 9). To design primers, we identified the most conserved regions of a Clustal-W multiple alignment within coding sequences of these genes (Fig. S1a-e). Primers were chosen according to consensus and degeneracy rules consistent with CODEHOP recommendations and predicted to have similar melting temperatures (Fig. 1a) (10). To validate these primers, we looked at four key properties. First, we determined primer specificity with capillary electrophoresis (Fig. 1b, S2a) and melt curve analysis (Fig. S2b), which demonstrated a single dominant amplicon for histones H1, H2A, H2B and H4. BLAST analysis of the dominant species confirmed histone isotype specific concordance across our primers (Fig. S2c). Degenerate primers for histone H3 produced the expected product, but demonstrated poor template-dependent amplification and so were not further pursued (Fig. S3a-g). Second, we quantified sensitivity across low template inputs with reproducible amplification at sub-nanogram cDNA levels (Fig. 1c, S2d). Third, we observed consistent and acceptable efficiencies by standard curve analysis across a 10,000-fold input range in two genetically distinct human cell lines (Fig. 1d). Fourth, we determined whether the assay captures dynamic changes in histone transcription following pharmacologic perturbation. Consistent with prior work showing reduction in RNA content in cells treated with aclarubicin (11), we observed a significant decrease in histone mRNA levels (Fig. 1e). We conclude that our assay reliably quantifies replication-coupled histone mRNA abundance and is responsive to modulating histone gene expression.

**Figure 1.**
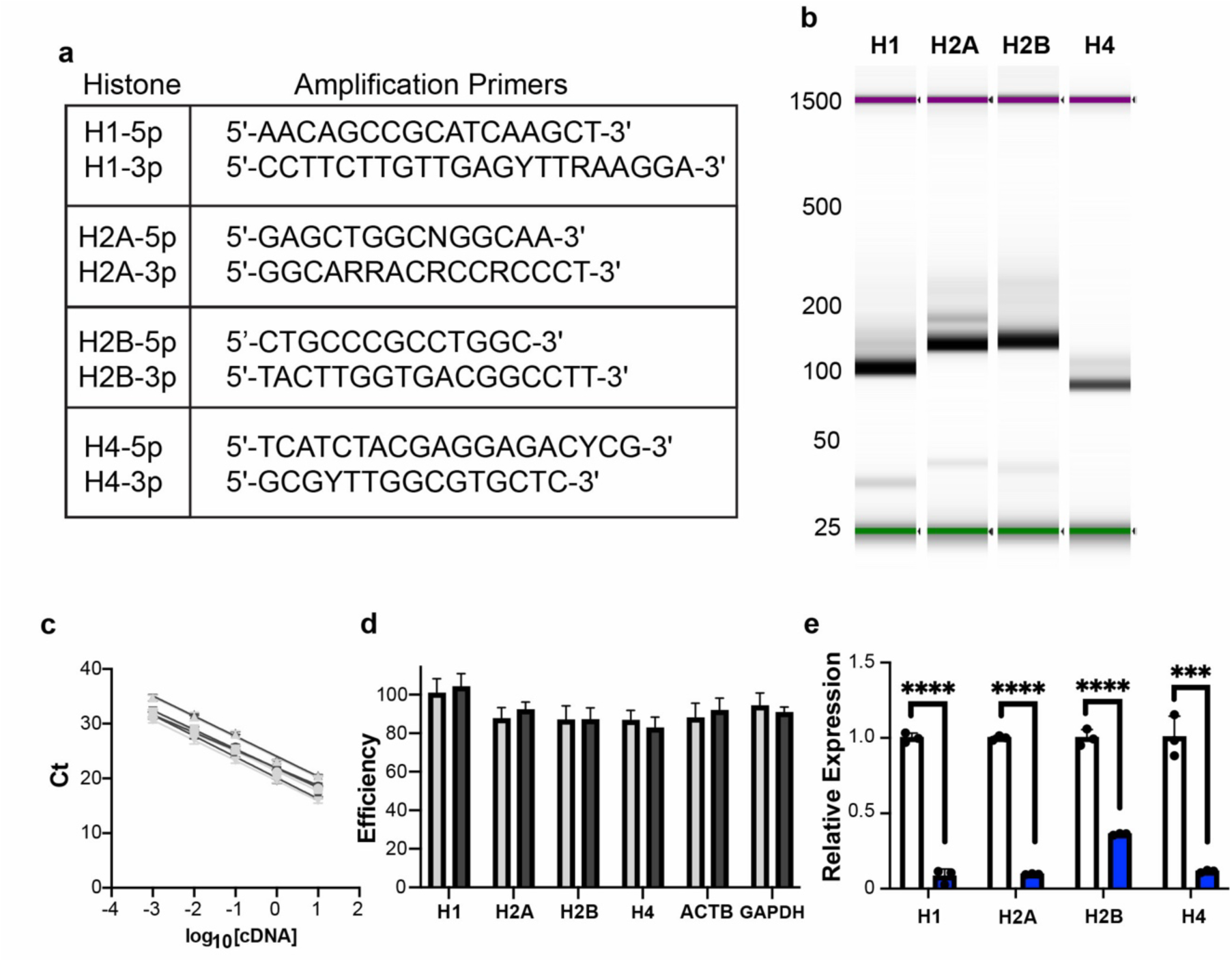
Degenerate PCR primers enable quantitative measurement of replication-coupled histone mRNA. **(a)** Primers targeting the most conserved regions of a Clustal-W alignment of histones H1, H2A, H2B, and H4. **(b)** Electropherogram of cDNA amplicons following PCR with bands of expected size confirm specificity. **(c)** Standard curves were generated using a serial dilution of cDNA following random hexamer primed reverse transcription from GM12878 (black) and K562 (gray) cells. Data points (H1 ●, H2A ▪, H2B ▴, H4 ▾) represent the mean (n=4 technical replicates; error bars indicate SD) and solid lines represent linear regression fits for each primer set. **(d)** Primer efficiency was calculated from the slope of each standard curve using E=10^−1/slope^−1 from (c). Bars represent mean efficiency, and error bars indicate 95% confidence intervals (CI) derived from the upper and lower limits of the regression slope for K562 (light gray) and GM12878 (dark gray). **(e)** Total histone mRNA following 4 h treatment in K562 cells with 100nM aclarubicin (blue) compared to DMSO controls (white) shows reduced histone levels. Bars represent mean ± SD of three biological replicates; points indicate individual replicates (n = 4 technical replicates). *** P < 0.001, **** P < 0.0001.

We applied our quantitative histone gene expression assay in a cancer versus non-cancer context using the CML-derived K562 and the immortalized GM12878 lymphoblastoid cell lines. Unlike other genes transcribed by RNAPII, the replication-coupled histone genes are restricted to S-phase to produce transcripts that are not polyadenylated, and instead undergo specialized 3’ stem-loop processing (5). We therefore performed reverse transcription using random hexamer primers and found a significantly increased amount of histone H1, H2A, H2B, and H4 transcripts in K562 compared to GM12878 (Fig. 2a). We next asked whether this increased histone output occurred in the context of cancer-associated hypertranscription. To compare transcriptional output, we measured total RNA and genomic DNA which revealed a greater RNA:gDNA ratio in the K562 leukemia line compared to the non-cancer GM12878 lymphoblastoid line (Fig. 2b). These findings demonstrate that increased replication-coupled histone mRNA abundance in K562 cells occurs in the setting of genome-wide hypertranscription.

**Figure 2:**
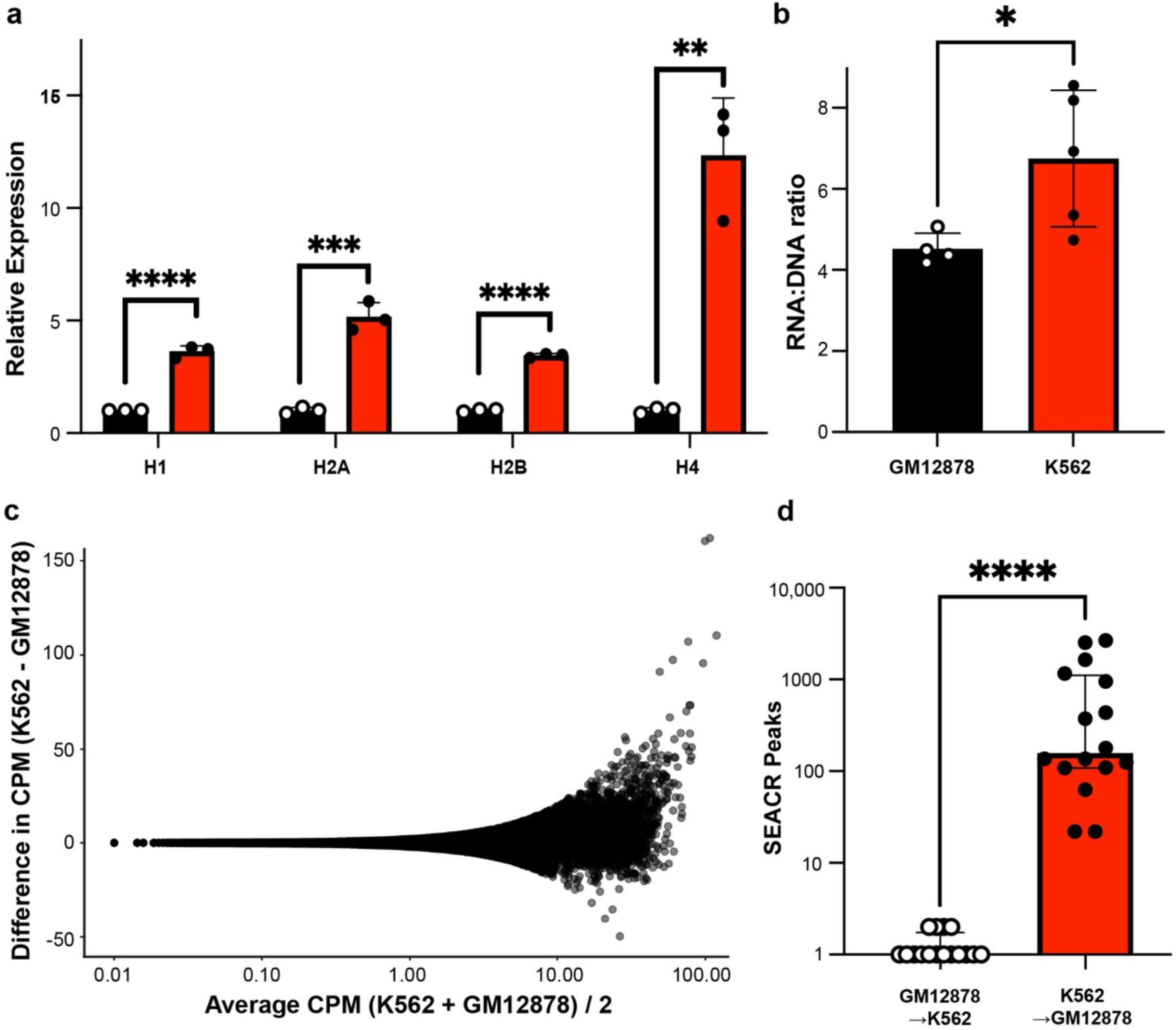
K562 cells exhibit histone overexpression and global hypertranscription compared to GM12878 cells. **(a)** Quantitative PCR following random hexamer priming for non-cancer GM12878 control (black) and cancer K562 (red) cells reveals increased total histone mRNA for H1, H2A, H2B, and H4. Bars represent mean ± SD of three biological replicates; points indicate individual replicates (n = 4 technical replicates). **P < 0.01; ***P < 0.001; ****P < 0.0001. **(b)** Direct quantification of total RNA and genomic DNA demonstrate K562 cells (n = 5) have a higher total RNA to gDNA ratio compared to GM12878 cells (n = 4). Bars represent mean ± SD. Statistical significance was assessed using a two-tailed unpaired t-test (P < 0.05). **(c)** RNAPII Ser5p CUTAC signal was quantified over ENCODE cCREs by counting overlapping fragments per cCRE and normalizing each library to counts per million mapped fragments (CPM; counts / total fragments × 10^6). For each cCRE, CPM values were averaged across K562 (n = 4) and GM12878 (n = 4) replicates. A Bland–Altman plot displays the per-cCRE difference in average CPM (K562 − GM12878) as a function of the average CPM across both cell lines (average of the two means; x-axis log-scaled with a +0.01 pseudocount), revealing a shift toward higher RNAPII Ser5p occupancy at many cCREs in K562 relative to GM12878, consistent with cancer-associated hypertranscription in the cancer cell line. **(d)** Reciprocal SEACR peak calling was performed for all pairwise comparisons between K562 and GM12878 datasets, alternately treating each sample as experimental versus control (n = 32 total comparisons). When K562 was used as the experimental sample against GM12878, hundreds to thousands of peaks were consistently identified (K562 → GM12878, median = 156.5), whereas the inverse comparison yielded no peaks in the majority of cases (GM12878 → K562, median = 0). Statistical testing of peak counts was used to summarize directional differences between comparisons (Mann–Whitney test, two-tailed, P < 0.0001) and displayed on a log10(count + 1) scale. Bars are median with IQR. Peaks were called using SEACR with norm and stringent arguments.

Since a majority of total RNA is ribosomal RNA, we also assessed genome-wide abundance of RNAPII, measured by CUTAC chromatin profiling. By profiling the initiating form of RNAPII (Pol II-Ser5p) in our cell line models, we found a genome-wide change across the ∼1 million candidate *cis*-regulatory elements (cCREs) defined by ENCODE (12), with more RNAPII in the K562 line than in the GM12878 line (Fig. 2c). As expected for hypertranscription, peak calling on RNAPII (1, 13) assigns many more transcribed sites in K562 cells (Fig. 2d).

Increased histone mRNA abundance might result from increased expression from some genes, or from increased expression of all genes. Total initiating RNAPII across the major and minor histone clusters was increased in K562 cells compared to GM12878 cells (Fig. 3a-b). However, some individual histone genes show increased or decreased amounts of RNAPII in K562 cells compared to the non-cancer control (Fig. 3c-h). We therefore tested whether individual replication-coupled histone genes are independently regulated within a set of primary human tumors. To do this, we analyzed samples with confirmed genome-wide hypertranscription with matched normal tissues from our prior study (1). We found that initiating RNAPII was elevated across the major and minor histone clusters overall, yet individual histone genes across all five histone types again exhibited discordant occupancy (Fig. S4a-b). These results show that replication-coupled histone genes can undergo gene-specific regulation of RNAPII activity within a genome-wide hypertranscription context.

**Figure 3:**
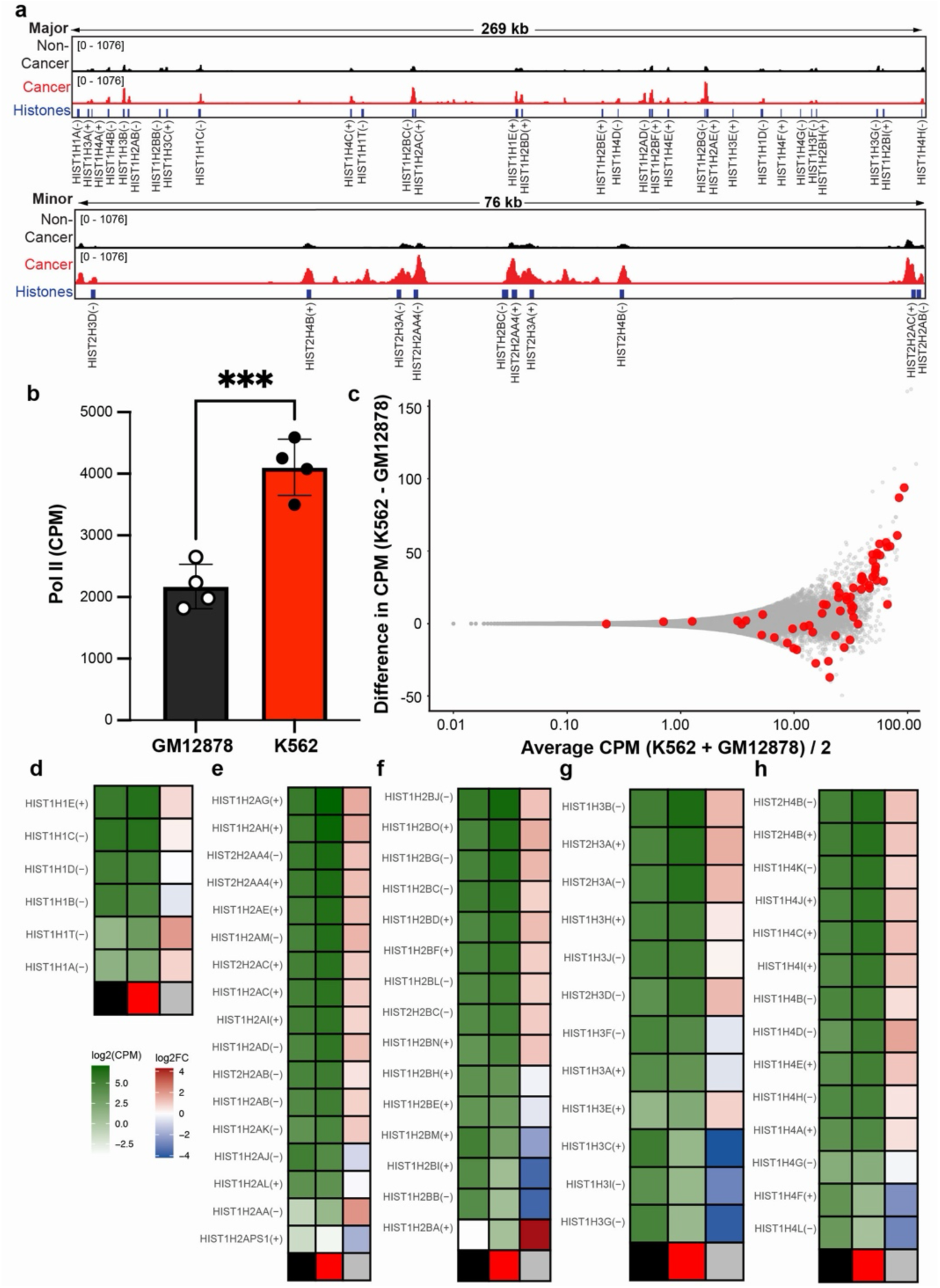
RNA RNAPII is elevated and heterogeneously distributed across the major and minor histone gene clusters. **(a**) Representative genome browser tracks are shown across the major and minor histone locus clusters for GM12878 (black) and K562 (red) cells with intervals for S-phase-dependent histone genes (blue). **(b)** Total HLB signal was quantified by counting CPM across the HLB interval. Each point represents one replicate (GM12878, n = 4; K562, n = 4). K562 exhibited higher HLB CPM than GM12878 (median 4163 vs 2109 CPM; ∼2.0-fold increase). Bars indicate mean with SD. One-tailed unpaired t-test, P = 0.0003. **(c)** Bland-Altman plot comparing RNAPII occupancy between K562 and GM12878 cells across cCRE and histone gene loci. Gray points represent ENCODE cCRE (same as in Fig. 2c); red points represent replication-coupled histone genes. **(d-h)** Heatmaps of RNAPII occupancy for individual histone genes within each histone (H1, H2A, H2B, H3, H4); see Supplemental Table 3 (1). Bottom rows indicate columns: black = GM12878, red = K562, Gray = log2 fold change (K562/GM12878). RNAPII levels are shown in a white-to-green gradient with log2 fold change displayed as blue-to-red gradient. Genes are ordered by descending order of RNAPII signal for K562.

We next performed reverse transcription with oligo-dT primers followed by quantitative PCR using our degenerate primers and found significantly greater levels of polyadenylated mRNA for histones H1, H2A, H2B, and H4 in K562 cells compared to GM12878 cells (Fig. 4a). We confirmed this increase using ENCODE-generated RNA-seq data from poly(A)-selected libraries and found significant enrichment of polyadenylated replication-coupled histone transcripts (2.9-fold; median 0.09% in GM12878 and 0.26% in K562; Fig. 4b) (14–16). As S-phase-dependent transcription of replication-coupled histone mRNAs generate a 3’ stem-loop product (5, 17, 18), this indicates altered histone mRNA processing in cancer cells.

**Figure 4:**
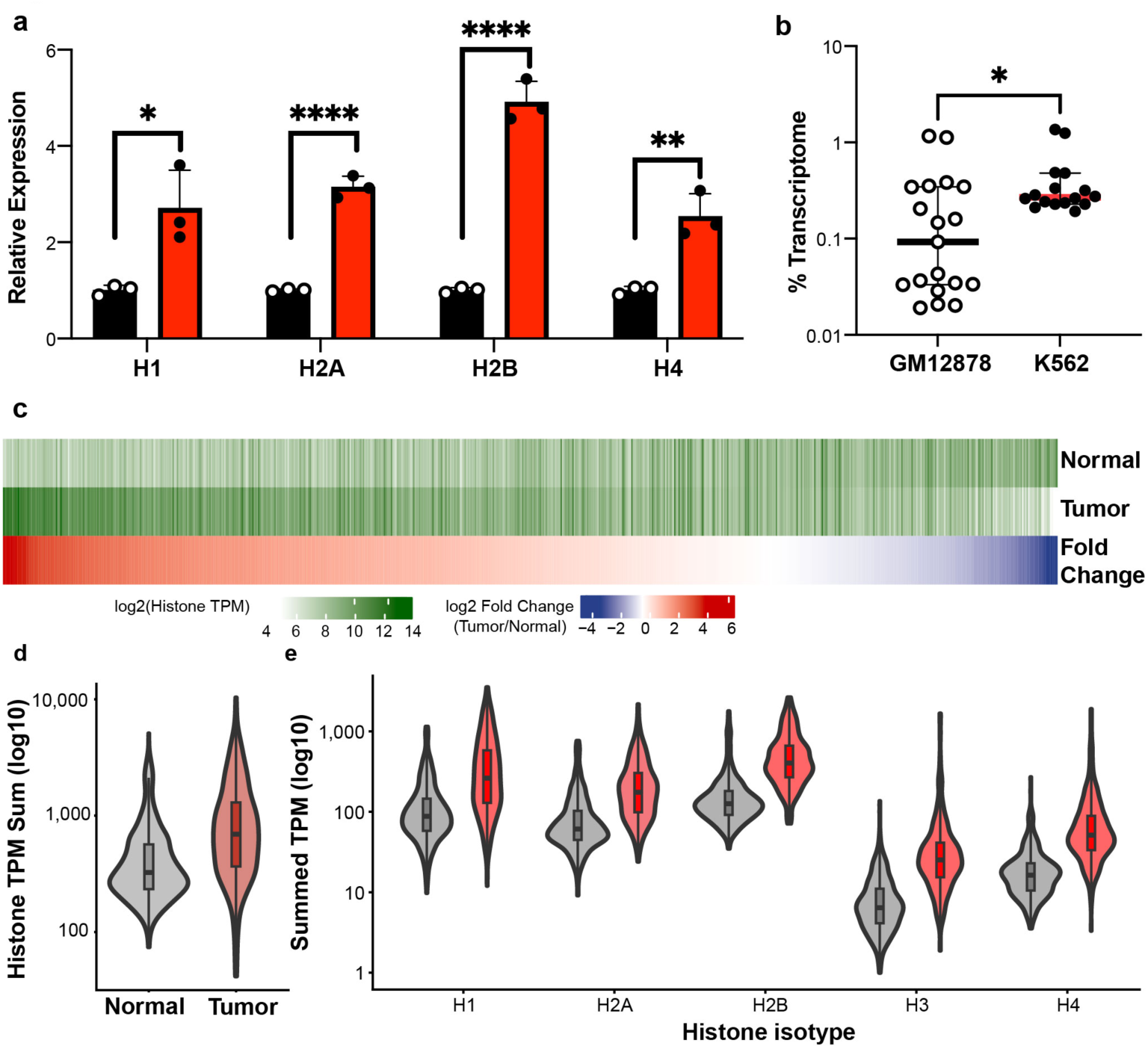
Polyadenylation of replication-coupled histone mRNA is widespread in human cancer. **(a)** Histone mRNA quantified by RT-qPCR using oligo-dT priming for GM12878 (black) and K562 (red) cells reveals increased amounts of polyadenylated histone mRNA for H1, H2A, H2B, and H4. Bars represent mean ± SD of three biological replicates; points indicate individual replicates (n = 4 technical replicates). *p < 0.05; **p < 0.01; ****p < 0.0001. **(b)** PolyA-selected RNA-seq for replication-coupled histone genes expressed as a proportion of total transcriptome (% transcriptome; log10 scale) in GM12878 and K562 cell lines shows significant increase in K562 compared to GM12878 cells. Each point is an isogenic replicate from ENCODE with median +/-95% CI (14–16). Mann-Whitney U test, P = 0.012. **(c)** Heatmap showing total polyadenylated replication-coupled histone gene expression across 715 matched tumor-normal pairs from 22 TCGA cancer types. Each column represents one patient. Top two rows show log2-transformed summed TPM of normal and tumor tissue, respectively (white to dark green). Bottom row shows tumor/normal log2 fold change color scaled symmetrically between the 1^st^ and 99^th^ percentile. **(d)** Polyadenylated histone gene expression for total replication-coupled histone genes is broadly upregulated in tumor relative to matched normal tissue across TCGA cancer types. Violin plots show the distribution of summed TPM across all replication-coupled histone genes in matched normal (gray) and primary tumor (red) samples from 715 patient pairs spanning 22 TCGA cancer types. Internal boxplots show the median and interquartile range. Tumor samples show a median 1.86-fold increase in histone TPM (72.9% of pairs upregulated; paired Wilcoxon signed-rank test, p = 2.07 x10^-49^). **(e)** Upregulation of polyadenylated histone mRNA is observed across all five histone isotypes. Violin plots show the distribution of summed TPM in matched normal (gray) and primary tumor (red) for each histone isotype across 521 patient pairs in which total polyadenylated histone TPM was higher in tumor than matched normal tissue. Boxplots show median and interquartile range. All five histone isotypes show significantly elevated tumor expression (paired Wilcoxon signed-rank test, BH-adjusted p < 9.29 x10^-69^).

ENCODE standardized RNA-seq data processing quantifies signal over all transcripts and does not distinguish whether these reads arise from polyadenylated or from non-polyadenylated transcripts. Canonical 3’ processing of replication-coupled histone genes depends on U7 snRNP-guided endonucleolytic cleavage by CPSF73 (19). We therefore inspected the distribution of RNA-seq coverage in ribosomal RNA depleted total RNA-seq and polyadenylated enriched RNA-seq libraries for K562 and GM12878 cells at the histone genes (Fig. S5a). H1-2, which can be expressed as a polyadenylated transcript secondary to cellular differentiation (20, 21) outside of S-phase (22), had the greatest difference in polyadenylated histone transcripts and demonstrated coverage that extended beyond the cleavage site that is unique to the polyadenylated isoform (Fig. S5b-e). Notably, a majority of the histone genes in K562 cells can be expressed as polyadenylated transcripts (Fig. S5f). Together, these data indicate these polyadenylated transcripts are produced from cryptic polyadenylation sites downstream of a subset of histone genes in human cancer.

To determine whether polyadenylated histone mRNA increases in cancer, we analyzed matched tumor-normal samples from The Cancer Genome Atlas (TCGA) that underwent polyadenylated RNA selection followed by sequencing (23). Across 715 patients spanning 22 cancer types, polyadenylated replication-coupled histone transcripts were elevated in tumor relative to matched normal tissue (Fig. 4c-d, S6a). Among the 521 patient tumors spanning 21 cancer types that exhibited greater polyadenylated histone levels compared to normal (Fig. S6b), all five histone isotypes were significantly elevated (Fig. 4e). These findings demonstrate that polyadenylation of replication-coupled histone mRNA is a widespread feature of human cancer.

In our quantitative PCR assay using degenerate histone gene primers, we noted that histone H2B had a greater relative expression of polyadenylated transcripts compared to histones H1, H2A, and H4 (Fig. 4a). To determine whether this preferential accumulation of polyadenylated H2B transcripts was a generalizable feature of human cancer, we calculated the tumor-normal difference for each isotype and ranked them within each of 521 TCGA patients with polyadenylated histone gene overexpression. Ranked by absolute tumor-normal difference, histone H2B was the most frequently top-ranked isotype, whereas histone H3 most frequently ranked last (Fig. 5a). We next tested whether this ranking was a consistent feature of polyadenylated histone abundance across cancer irrespective of matched normal changes. To determine the tumor intrinsic relative abundance of each histone isotype, we calculated the difference between each histone and the mean of the remaining four histones. In 10,765 cancer samples across 33 cancer types, we again find a histone H2B-first and histone H3-last trend (Fig. 5b-c, S7a-b). Notably, this pattern was recapitulated across seven cancer transcriptional subtypes despite substantial differences in global transcriptional programs and clinical behaviors (Fig. 5b-c). We suggest a conserved hierarchy of histone isotype abundance in cancer associated hypertranscription that is affected by baseline expression levels of each histone type.

**Figure 5:**
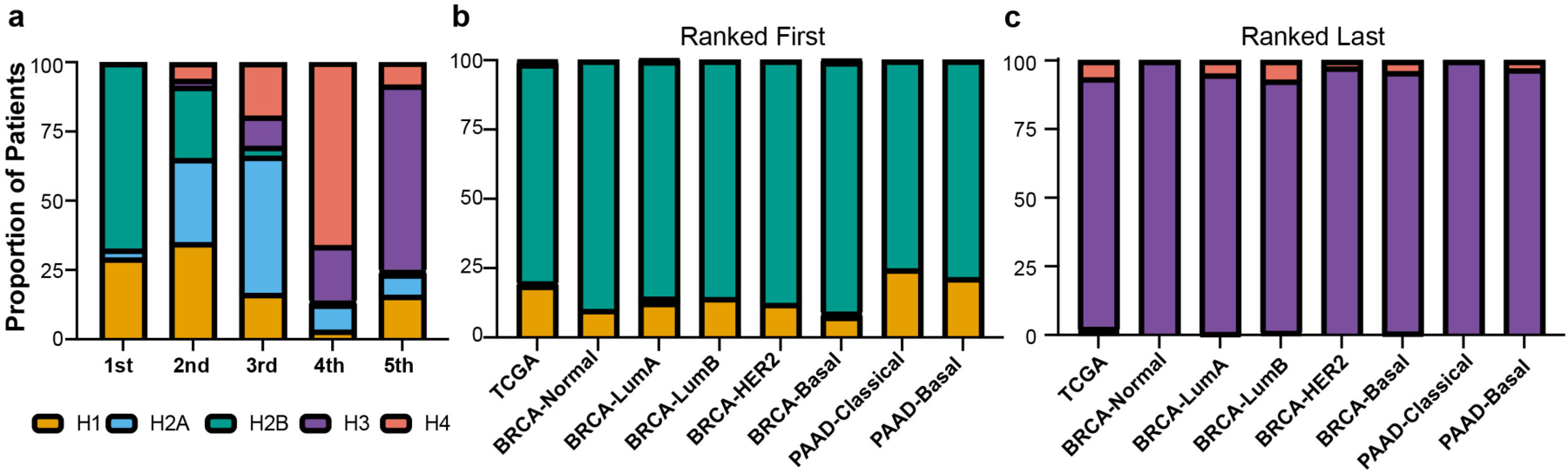
Polyadenylated histone mRNA accumulation in cancer is dominated by histone H2B transcripts across diverse tumor types and molecular subtypes. **(a)** Proportion of histone-elevated tumor-normal pairs at each rank position for each histone isotype, ranked by absolute increase in tumor minus normal, where 1st indicates the largest increase. Histone H2B ranked first in the majority of patients (67.4%), while histone H3 ranked last in the majority of patients (66.6%) in cancer samples with elevated total histone abundance (n = 521 pairs across 22 cancer types). (B, C) For each tumor sample, the relative abundance of each histone isotype was assessed by calculating the difference between a given isotype and the mean of the remaining four isotypes. Samples ranked first **(b)** have the highest positive difference, indicating that isotype is more abundant relative to the others. Conversely, samples ranked last **(c)** have the lowest (most negative) difference, indicating that isotype is the least abundant relative to the others. The proportion of each histone isotype ranking first and last was determined across pan-cancer TCGA matched tumor samples, breast cancer (BRCA) PAM50 subtypes and pancreas cancer (PAAD) Moffitt molecular subtypes. Histone H2B ranked first in the majority of samples across all groups (range: 75–90%), and histone H3 ranked last in the vast majority of samples across all groups (range: 92–100%). TCGA, n = 10,765 cancer samples; BRCA Normal n=40, LumA n=571, LumB n=209, Her2 n=82, Basal n=197; PAAD Classical n=85, Basal n=65.

The replication-coupled histone genes generate transcripts during S-phase by coordinating processing factors to histone locus bodies (HLBs), a subnuclear structure that contains all the components needed for histone gene transcription and transcript processing (24). RNAPII is an integral component of the HLB and accumulates during both S- and gap-phases of the cell cycle (25, 26). Nascent transcription of the histone genes occurs at the HLB throughout the cell cycle in a context-specific manner in Drosophila (25). Similarly, the non-cancer GM12878 human cell line accumulates initiating RNAPII within HLBs mainly during S-phase, but initiating RNAPII remains associated with histone genes throughout interphase in K562 cells (26). To examine the abundance of polyadenylated histone transcripts throughout the cell-cycle, we performed fluorescence-activated cell sorting to isolate populations of K562 cells in G1-, S-, and G2-phases. Cell cycle phase-specific quantification of polyadenylated histone mRNAs revealed an abundance of these transcripts throughout the cell cycle (Fig 6a-b). We find that these transcripts are proportionally more abundant in gap phases of K562 cells.

**Figure 6.**
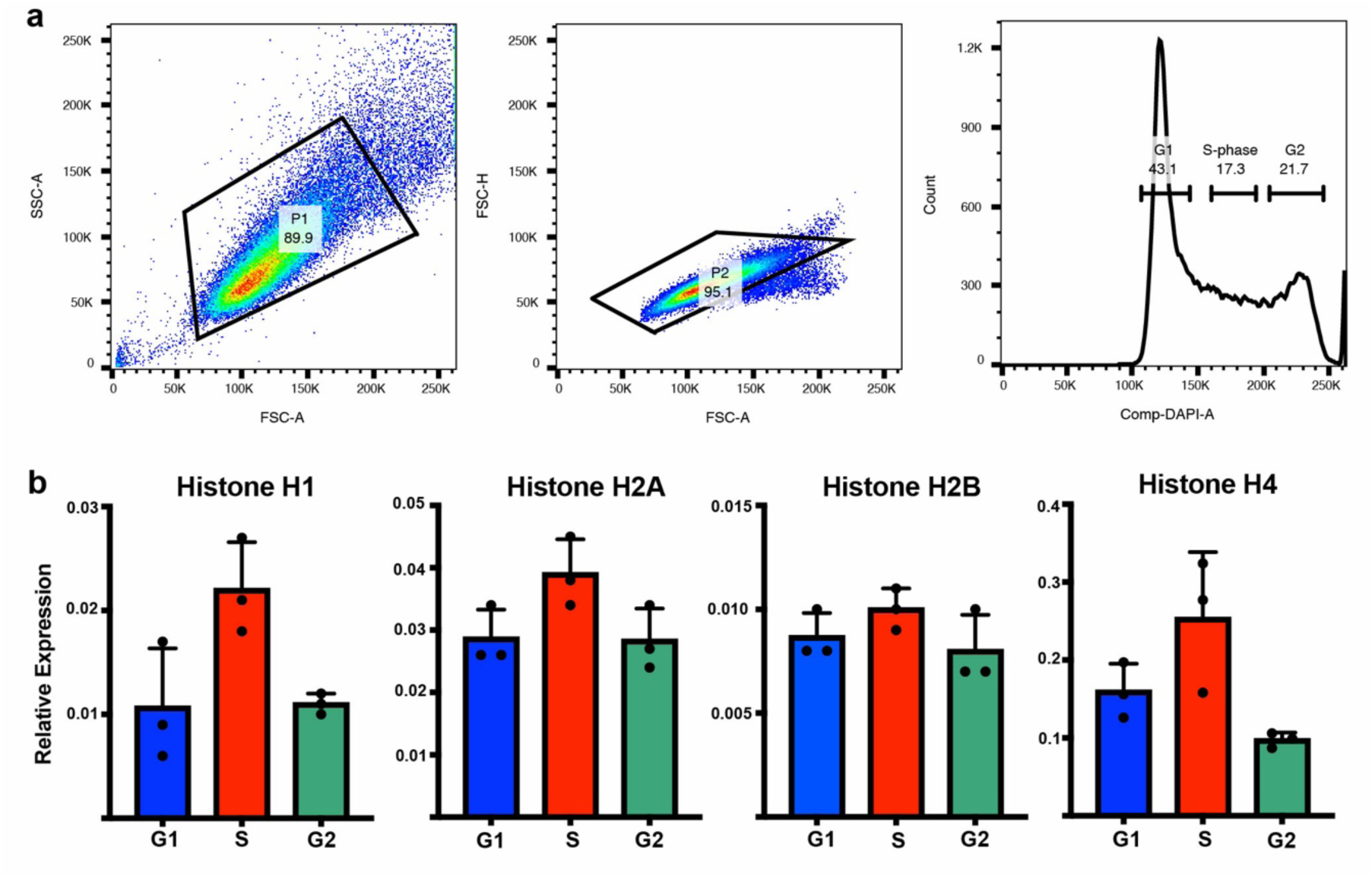
Quantification of polyadenylated histone transcripts across cell cycle phases. **(a-b)** Following DAPI staining, K562 cells were collected by fluorescence-activated cell sorting into representative G1 (blue), S (red), and G2 (green) populations based on DNA content. RNA was purified from each sorted population, and cDNA was synthesized by oligo(dT)-primed reverse transcription. RT-qPCR was performed in technical quadruplicate for each biological replicate (n=3). ΔCt was calculated for each histone gene as the difference between the histone Ct value and the geometric mean of the ACTB and GAPDH Ct values. Relative expression was calculated as 2^−ΔCt for each histone.

We next investigated the regulatory mechanisms that govern the abundance of polyadenylated histone mRNAs by leveraging publicly available RNA-seq datasets (Supplemental Table 2). Because replication-coupled histone gene expression during S-phase requires coordinated transcription, we hypothesized that perturbation of RNAPII regulatory factors could alter polyadenylated histone mRNA abundance. Prior work showed that inhibition of CDK7 – a kinase that promotes RNAPII promoter escape by releasing initiation factors and Mediator (27) – increased polyadenylated histone mRNA (Fig. 7a, S8a-b) (28, 29). Since RNAPII pausing is downstream of promoter escape (30), we investigated whether disruption of P-TEFb mediated pause-release had similar effects. Indeed, another study that selectively inhibited and degraded CDK9, the catalytic subunit of P-TEFb, increased polyadenylated histone mRNA (Fig. 7b) (31). We observed a similar increase in additional published datasets following selective inhibition and degradation of BRD4, which directly interacts with P-TEFb (Fig. 7c, S8c-g) (32, 33). In contrast, prior work investigating decreased CDK12/13 activity, which regulates RNAPII elongation, had minimal effects in this one dataset (Fig. 7a, S8a) (28). We further evaluated if Integrator-PP2A-mediated dephosphorylation of the RNAPII CTD (34, 35), and found that its loss did not significantly change polyadenylated histone transcript levels (Fig. 7d) (36).

**Figure 7:**
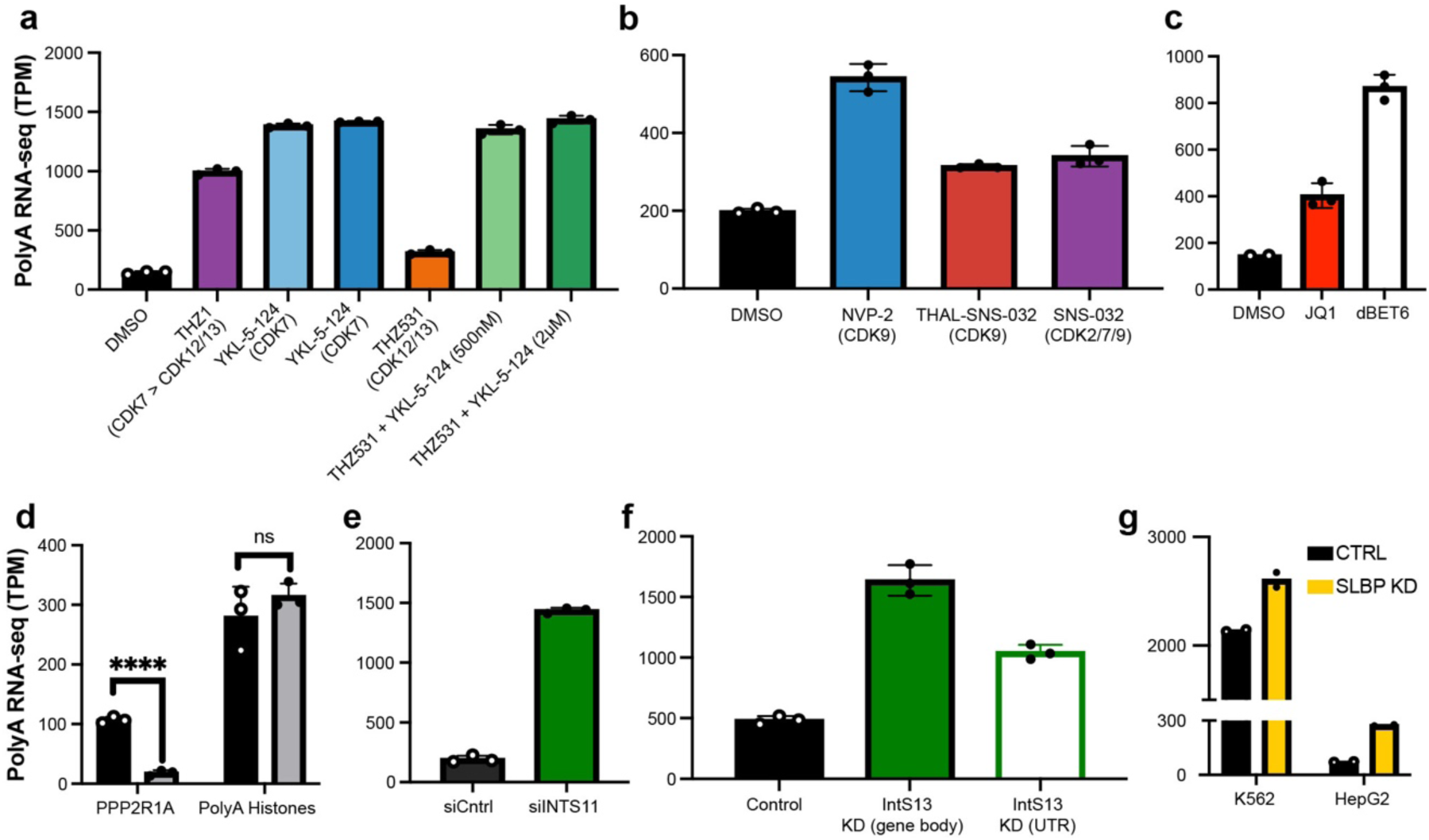
Perturbation of promoter-proximal RNAPII activity increases polyadenylated histone gene expression. **(a)** HAP1 cells treated for 24 hours with DMSO, YKL-5-124 (CDK7-selective), THZ1 (CDK7/12/13), THZ531 (CDK12/13), or THZ531 combined with YKL-5-124 (28). One-way ANOVA with Dunnett’s test; all comparisons significant (P adj < 0.0001). **(b)** MOLT4 cells treated for 6 hours with the selective CDK9 inhibitor NVP-2, the selective CDK9 degrader THAL-SNS-032, or the multi-target CDK2/7/9 inhibitor SNS-032 (31). One-way ANOVA with Dunnett’s test; all comparisons significant (P adj ≤ 0.0006). **(c)** MOLT4 cells treated with dBET6 or JQ1 for 6 hours (32). One-way ANOVA with Dunnett’s test; all comparisons significant (P adj ≤ 0.0034) following Grubbs outlier test. **(d)** HeLa cells following PP2A (PP2A-A) knockdown versus control (36). Unpaired t-test, ***P < 0.0001, ns = not significant. **(e)** HeLa-LTR-Luciferase cells transfected with siRNA targeting Integrator subunit 11 (INTS11) or scrambled control (38). Unpaired two-tailed t-test, P <0.0001. **(f)** ARPE-19 cells with control knockdown, INTS13 coding sequence knockdown, or INTS13 UTR-targeting knockdown (63). One-way ANOVA with Dunnett’s test; all comparisons significant, P adj ≤ 0.0003. **(g)** PolyA histone transcript levels in K562 and HepG2 cells following SLBP knockdown (ENCODE) (14–16).

Beyond RNAPII regulation, we considered whether spatial localization of RNAPII at HLBs contributed to polyadenylated histone transcript levels. We found that published knockdown of INTS11 increased levels of polyadenylated histone mRNA (Fig. 7e) (38), consistent with increased polyadenylated transcripts of four individual histone genes following INTS9 depletion (39). Because S-phase-dependent histone transcription generates a conserved 3’ stem-loop pre-mRNA for processing, we investigated a prior study that characterized the effects following perturbation of the functional module of Integrator (INTS10-INTS13-INTS14) that preferentially binds stem-loop RNA (40). In these data, knockdown of INTS13 produced a significant increase in polyadenylated histone transcripts (Fig. 7f). Since stem-loop binding protein (SLBP) is required for U7 snRNP mediated cleavage of the 3’ stem-loop, SLBP loss redirects individual histone loci to generate polyadenylated transcripts (41). Similarly, data from ENCODE demonstrated that SLBP depletion elevated total polyadenylated histone mRNA (Fig. 7g) (14–16).

The selective increases in polyadenylated histone transcripts following diminished RNAPII initiation or pause-release suggests that promoter-proximal RNAPII regulation is required to maintain the fidelity of replication-coupled histone mRNA processing. This is consistent with the short coding sequences of the replication-coupled histone genes (median 390 bp), which are less dependent on CDK12/13-mediated processive elongation and 3′-end maturation of long, multi-exon transcripts (42). The lack of response following decreased INTAC-mediated CTD phosphatase activity, despite its role in regulating RNAPII pause-release to attenuate transcription (34, 35), further indicates that CTD phosphorylation state is not the critical factor of polyadenylated histone transcript levels. Instead, Integrator subunits and stem-loop processing factors at the HLB (41) are key determinants coupling S-phase transcription to stem-loop-dependent 3’ processing, as loss of either increase polyadenylated histone transcripts. Factors involved in transcription elongation (43–47) and histone pre-mRNA processing (48) are required for proper replication-coupled histone mRNA 3’-end formation. Loss of these factors disrupts precise stem-loop cleavage, allowing RNAPII procession beyond the normal processing site, and leads to the accumulation of aberrant polyadenylated histone transcripts. Our findings suggest a model in which canonical S-phase-dependent histone mRNA processing depends on adequately coordinated RNAPII occupancy at the HLB, and its disruption results in altered processing efficiency or changes in mRNA stability which redirects histone gene output toward unscheduled, polyadenylated transcripts.

Our study also demonstrates that polyadenylated replication-coupled histone mRNAs accumulate pervasively across human cancers. The capacity for HLB-coordinated stem-loop processing during S-phase may be limiting relative to the elevated transcriptional demand imposed by cancer-associated hypertranscription, such that a fraction of histone transcripts escape S-phase-coupled processing and accumulate as polyadenylated mRNAs. The conserved enrichment of H2B across diverse cancer types and transcriptional subtypes is consistent with this model, as H2B lacks replication-independent variants that could otherwise fulfill its structural role outside of S-phase. Conversely, the comparatively modest enrichment of histone H3 likely reflects the contributions of replacement histone H3.3 to nucleosome deposition (7, 49), suggesting the presence of a feedback mechanisms responsive to histone isotype-specific abundance. This preferential enrichment may also reflect differential promoter architecture, as factors such as OTF1 (50) stimulate histone H2B gene transcription. Our finding that cancer cells accumulate polyadenylated isoforms of the replication-coupled histone genes throughout the cell cycle resembles how cells process the mRNA for the histone variant H2AX. The H2AX gene generates a stem-loop processed transcript isoform in S-phase cells, while polyadenylated transcripts are expressed throughout the cell cycle (51, 52). Additionally, histones H3.3, H2A.Z and CENP-A are variant histones that have polyadenylated tails and are available for deposition throughout the cell cycle (53). We propose that cancer-associated hypertranscription during gap phase of the cell cycle may drive unscheduled replication-coupled histone gene expression, resulting in the accumulation of polyadenylated histone mRNA.

These alternatively processed histone transcripts provide a reservoir of stable mRNAs with four complementary consequences in cancer. First, elevated histone synthesis due to greater transcript levels would support chromatin packaging during proliferation in cancer cells compared to normal cells, providing an intrinsic unifying mechanism for proliferative Hallmarks of Cancer (1, 54). Second, an abundance of stable histone mRNA outside of S-phase may compress the G1- to-S transition, insofar as histones are rate limiting (6, 55), consistent with evidence that cell-cycle duration is the key determinant of oncogenic transformation (56). Third, unscheduled replication-coupled histone H3 synthesis during gap phase may generate polyadenylated transcripts that ultimately compete with the centromeric variant CENP-A to promote chromosomal instability and provide the genetic substrate for cancer initiation (1, 4, 57, 58). Fourth, terminally differentiated mammalian cells express polyadenylated histone transcripts from a subset of genes in the major and minor histone clusters, many of which encode histone H2B, and likely function in normal tissues to replace histones that are damaged (22). Disseminated cancer cells remain quiescent prior to metastasis (59), and these polyadenylated histone transcripts may then provide an additional buffer to maintain the chromatin landscape in their dormant state. Whether these transcripts functionally contribute to each of these processes to influence clinical outcomes in cancer remain to be determined.

## Materials and Methods

### Cell culture

Human K562 chronic myelogenous leukemia cells (American Type Culture Collection (ATCC) cat. no. CCL-243) were cultured in liquid suspension in IMDM (ATCC) with 10% HI-FBS (Hyclone SH3007003). GM12878 lymphoblastoid cells from Coriell Institute were grown in RPMI (Gibco #11875-093) with 15% HI-FBS (Seradigm 1500-500H). Cell lines were authenticated by short tandem repeat profiling and confirmed mycoplasma-free. Cells in log phase were collected by centrifugation and washed in 1xPBS prior to pellets being flash frozen in liquid N2 and stored at -70°C.

### Nucleic acid isolation

Total RNA and genomic DNA were extracted following Macherey-Nagel NucleoSpin® RNA and DNA Buffer protocols. RNA quality and integrity were assessed via fragment analysis with the Agilent 4200 Tapestation on a HSRNA screentape. Rnasin® Rnase Inhibitor (Promega) was added to the purified RNA following product recommendations (1 U/µL) and stored at -20°C short-term or -70°C for longer. Nucleic acids were quantified by spectrophotometry.

### cDNA synthesis

Total RNA was transcribed into cDNA following the Moloney Murine Leukemia Virus Reverse Transcriptase Kit protocol (M-MLV RT, Invitrogen). Random hexamer and oligo-dT synthesized cDNA were prepared in a 20 µL reaction by mixing 10 µL of RNA (1 µg of RNA/reaction), 1 µL of random hexamers (2.5 µM final concentration) or oligo-dT primers (25 ng/µL final), and 1µL of dNTP mix (0.5 mM final). This mixture was heated to 65°C for 5 minutes then cooled on ice before adding 4 µL 5X First-Strand Buffer (1X final), 2 µL 100 mM DTT (10 mM final), and 1 µL 40U/µL Rnasin® Rnase Inhibitor (2 U/µL final). It was then heated for 2 minutes at 37°C before the addition of 1 µL of 200 U/µL M-MLV Reverse Transcriptase (10U/µL final). Both mixtures, regardless of synthesis primer, were heated for 10 minutes at 25°C then 50 minutes at 37°C. The transcriptase was inactivated by incubation at 70°C for 15 minutes and residual RNA digested by addition of 0.5 µL RNase H (0.1U/uL) at 37°C for 20 minutes. Reactions were then stored at -20°C.

### qPCR

cDNA levels were determined by PCR using the 384-well ABI QuantStudio™ 5 RealTime PCR System. Reactions were prepared in 384-well plates with each reaction containing 10 µL of 2x SYBR™ Green PCR Master Mix (Applied Biosystems, 1x final), 3.2 µL of a primer mix containing the forward and reverse primers at 10 µM each (1.6 µM each final), 2 µL of cDNA, and nuclease-free water to a final volume of 20 µL. Annealing temperature were empirically determined by comparing Ct values across a range. Reactions were incubated at 55°C 2 minutes, 95°C 5 minutes, then 40 cycles of 95°C for 15 seconds and 58°C for 30 seconds, followed by 40°C for 1 minute then a melt curve from 95°C to 60°C for 1 minute at each temperature; ramp rate throughout the program was 1.6 C/s. Ct and derivative melt-curve values were automatically calculated using QuantStudio™ Design and Analysis Software V1.4.3. After cycling, PCR product was visualized using the Agilent 4200 Tapestation with an HSD1000 screentape. Amplicons were purified using the Macherey-Nagel Nucleospin Gel and PCR Cleanup Kit (740609.50). Following purification, Premium PCR Sequencing was performed by Plasmidsaurus using Oxford Nanopore Technology with custom analysis and annotation (Plasmidsaurus, San Francisco, CA).

H1-5p: 5’-AACAGCCGCATCAAGCT-3’
H1-3p: 5’-CCTTCTTGTTGAGYTTRAAGGA-3’
H2A-5p: 5’-GAGCTGGCNGGCAA-3’
H2A-3p: 5’-GGCARRACRCCRCCCT-3’
H2B-5p: 5’-CTGCCCGCCTGGC-3’
H2B-3p: 5’-TACTTGGTGACGGCCTT-3’
H3-5p: 5’-GTGATGGCGCTGCA-3’
H3-3p: 5’-CCACGGATGCGGCG-3’
H4-5p: 5’-TCATCTACGAGGAGACYCG-3’
H4-3p: 5’-GCGYTTGGCGTGCTC-3’
hGAPDH_F: 5’-GTCTCCTCTGACTTCAACAGCG-3’
hGAPDH_R: 5’-ACCACCCTGTTGCTGTAGCCAA-3’
ACTB_fwd: 5’-CACCATTGGCAATGAGCGGTTC-3’
ACTB_rev: 5’-AGGTCTTTGCGGATGTCCACGT-3’

### CUTAC-direct for whole cells

CUTAC was performed as previously described (60), with some modifications. Briefly, cells were mixed with activated Concanavalin A beads and resuspended in Triton-wash buffer (20 mM HEPES pH 7.5, 150 mM NaCl, 0.5 mM spermidine, 0.05% Triton-X100 and Roche EDTA-free protease inhibitor). After successive incubations in Wash buffer for ≥1 hour in primary antibody (RNAPII-Ser5p: Cell Signaling Technologies cat. no. 13523, lot 3) and then 1 hour in secondary antibody (Guinea pig α-rabbit antibody: Antibodies online cat. no. ABIN101961, lot 46671), the beads were washed and resuspended in pAG-Tn5 preloaded with mosaic end adapters (Epicypher cat. no. 15-1117, 1:20) in Triton-wash buffer for 1 hour. Incubations were done at room temperature in 25 µL volumes in PCR tubes. Tagmentation was performed for 1 hour in 10 mM TAPS pH 8.5, 20% N,N-dimethylformamide, 5 mM MgCl_2 a_t 55 °C. Fragment release was performed in 5 µL 1% SDS supplemented with 1:10 Thermolabile Proteinase K (New England Biolabs cat. no. P8111S) at 37 °C 1 hour followed by 58 °C 1 hour. SDS was quenched by addition of 15 µL 6% Triton-X100 and PCR was performed by addition of 2 µL each barcoded 10 mM i5 and i7 primer solutions and 25 µL NEBNext 2X PCR Master mix (New England Biolabs cat. no. ME541L). The following cycling conditions were used: Cycle 1: 58 °C for 5 minutes; Cycle 2: 72 °C for 5 minutes; Cycle 3: 98 °C for 5 minutes; Cycle 4: 98 °C for 10 seconds; Cycle 5: 60 °C for 10 seconds; Repeat Cycles 4-5 11 times; 72 °C for 1 minutes; Hold at 8 °C. Clean-up was performed using HighPrep PCR Cleanup beads (Magbio Genomics cat. no. AC-60500) following manufacturer’s instructions. A detailed step-by-step protocol is available at Protocols.io: https://www.protocols.io/view/cut-amp-tag-direct-for-whole-cells-with-cutac-x54v9mkmzg3e/v5

### Fluorescence-Activated Cell Sorting

To purify G1, S, G2 population of K562 cells, 10 million cells were washed in 1 ml PBS and resuspended in 100 µl of 3% glyoxal fixation solution (61) with 1:25 RNase Inhibitor (RNasin Plus; Promega N261B) prior to incubation for 15 minutes on ice. 1 ml of PBS with 1:100 RNase Inhibitor was added and cells were centrifuged at 2000 x g for 3 minutes at 4 °C. Cells were resuspended in 100 µl of ice-cold methanol (100%) with 1:25 RNase inhibitor and stored at -20 °C. Prior to sorting, all samples were filtered and labeled with DAPI (Sigma, D9542) at a final concentration 5µg/mL. Cells were collected in 1.5 mL RNase-free tubes pre-coated with 1% BSA in PBS supplemented with 1:25 RNase Inhibitor. Three cell cycle fractions were isolated on a BD FACSymphony S6 Cell Sorter based on DAPI intensity. Pellets were flash frozen in liquid N2 and stored at -70°C.

### DNA sequencing and data processing

The size distributions and molar concentration of libraries were determined using an Agilent 4200 TapeStation. Up to 96 barcoded libraries were pooled at approximately equimolar concentration for sequencing. Paired-end 50x50 bp sequencing on the Illumina NextSeq 2000 platform was performed by the Fred Hutchinson Cancer Center Genomics Shared Resources. This yielded 5–20 million reads per antibody. Adapters were clipped by cutadapt version 4.4 with parameters --nextseq-trim 20 -m 20 -a AGATCGGAAGAGCACACGTCTGAACTCCAGTCA -A AGATCGGAAGAGCGTCGTGTAGGGAAAGAGTGT -Z

Clipped reads were aligned by Bowtie2 version 2.5.1 to the *Homo sapiens* hg19 reference sequences from UCSC with parameters --very-sensitive-local --soft-clipped-unmapped-tlen -- dovetail --no-mixed --no-discordant -q --phred33 -I 10 -X 1000

### Data analysis

For quantitative PCR, Ct values were normalized to the geometric mean of GAPDH and ACTB to generate ΔCt values. Relative expression plotted as 2^^−ΔΔCt^. Statistical significance was assessed using unpaired two-tailed t-tests.

For chromatin profiling, properly paired reads were extracted from the alignments by samtools bamtobed command into mapped fragment bed files.

RNAPII Ser5p CUTAC signal was quantified over ENCODE cCREs by counting overlapping fragments per cCRE (bedtools intersect -c) and normalizing each library to counts per million mapped fragments (CPM; counts / total fragments × 10^6^). For each cCRE, CPM values were averaged across K562 and GM12878 replicates. Prior to reciprocal peak calling, fragments falling within RepeatMasker-annotated repetitive elements from UCSC were discarded using bedtools intersect -v. BedGraphs were generated using bedtools genomecov -bg. Peaks were called using SEACR with norm and stringent arguments. Human replication-coupled histone genes were annotated as previously described (1).

Normalized count tracks were visualized in IGV and have the fraction of counts at each base pair multiplied by the length of the genome so that if the counts were uniformly distributed there would be 1 at each base pair.

For major and minor histone gene annotation for transcriptomic analysis, we downloaded the HUGO Gene Nomenclature Committee (HGNC) defined gene group, “Histones” and selected genes that fell within 6p22 or 1q21.

We downloaded gene expression quantification files from RNA-seq generated from polyA-selected libraries from ENCODE (14–16) using experiment and accession identifiers in Supplemental Table 1. Gene-level quantifications (TPM) were generated from ENCODE processed data (GENCODE v29 annotation on the GRCh38 human reference genome). Histone genes were quantified by cross-referencing gene IDs against a histone gene annotation from the HUGO Gene Nomenclature Committee (HGNC). Histone-specific TPM values were extracted across all samples and percent transcriptome was quantified by calculating the aggregate histone TPM as a proportion of total sample TPM.

For TCGA analysis for matched tumor-normals, we used TCGAbiolinks (62) and first identified 22 of 33 TCGA projects containing at least one matched tumor-normal pair and downloaded gene expression quantification data for all primary tumor and solid tissue normal samples from those projects. RNA-seq data were retrieved for 22 cancer types and analysis was restricted to patients with both a primary tumor and a matched normal sample were retained, yielding 715 paired samples across all cancer types. For each patient, TPM values from all replication-coupled histone genes (major and minor cluster genes from our curated list) were summed per sample using HGNC IDs as gene identifiers, and tumor-to-normal fold change was calculated as the ratio of tumor to normal summed histone TPM. Differential expression was assessed per patient using a paired Wilcoxon signed-rank test. Per-cancer-type breakdowns of upregulated versus downregulated patients were summarized across all 22 cohorts.

For our TCGA analysis of a dominant histone isotype, we first identified a cohort of patients with elevated total histone expression (tumor TPM sum > normal TPM sum) yielding 521 patients. For each patient and histone isotype (H1, H2A, H2B, H3, H4), we summed TPM values across individual histone genes within the isotype for tumor and normal samples separately. We calculated the tumor-normal difference for each isotype per patient, then rank-ordered the five isotypes by magnitude of the difference. We then computed the proportion of patients for which each isotype ranked as having the largest absolute change.

For our histone isotype tumor intrinsic analysis, per-isotype histone TPM was summed across all replication-coupled genes within each isotype class (H1, H2A, H2B, H3, H4). For each sample, a leave-one-out contrast was computed by subtracting the mean TPM of the remaining four isotypes from each histone isotype of interest. Isotypes were then ranked within each sample from highest to lowest contrast, and the frequency with which each isotype ranked first (most relatively abundant) or last (least relatively abundant) was tallied across samples. This analysis was performed on pan-cancer TCGA samples (n=10,765), BRCA primary tumors stratified by PAM50 molecular subtype (n=1,099), and PAAD primary tumors stratified by Moffitt transcriptional subtype (n=150).

For RNA-seq from Gene Expression Omnibus, we identified publicly available RNA-seq datasets from GEO that used poly(A)-selected library preparation (Supplemental Table 2). For each dataset, we reviewed the associated methods and GEO metadata to confirm library preparation strategy and experimental design. We then downloaded the NCBI-generated normalized counts matrices from each Series and filtered for histone genes by NCBI Gene ID using our curated histone gene table derived from the HGNC “Histones” gene group. RNA-seq TPM was plotted mean and error bars indicate SD. Total histone TPM was computed per sample by summing the expression values of all matched histone genes across biological replicates. GEO series and samples are located in Supplemental Table 2.

CDS lengths were calculated from the RefSeq GRCh38 GTF annotation by summing CDS intervals per NCBI GeneID and intersecting with the curated histone gene set.

## Acknowledgments

We thank Christine Codomo and Doris Xu for technical assistance; the Fred Hutch Genomics Shared Resource for sequencing and data processing; Erik Huynh in the Flow Cytometry Shared Resource for assistance with cell sorting; and the ENCODE Consortium and the ENCODE production laboratories for generating the datasets used in this study. The results published here are in whole or part based upon data generated by the TCGA Research Network: https://www.cancer.gov/tcga. This work was supported by the Howard Hughes Medical Institute (S.H.) and grant # K12CA076930 from the National Cancer Institute (R.M.P.).

## Author contributions

R.M.P., K.A., and S.H. conceived the study. R.M.P., T.B., G.R.R., and K.K.N. performed the experiments. R.M.P. analyzed the data and wrote the original draft. K.A. and S.H. supervised the study and reviewed and edited the manuscript. All authors reviewed and approved the final manuscript.

## Data Availability

Data generated in this study have been deposited at the Gene Expression Omnibus: GSE327162

---

**Supplemental Figure 1.**
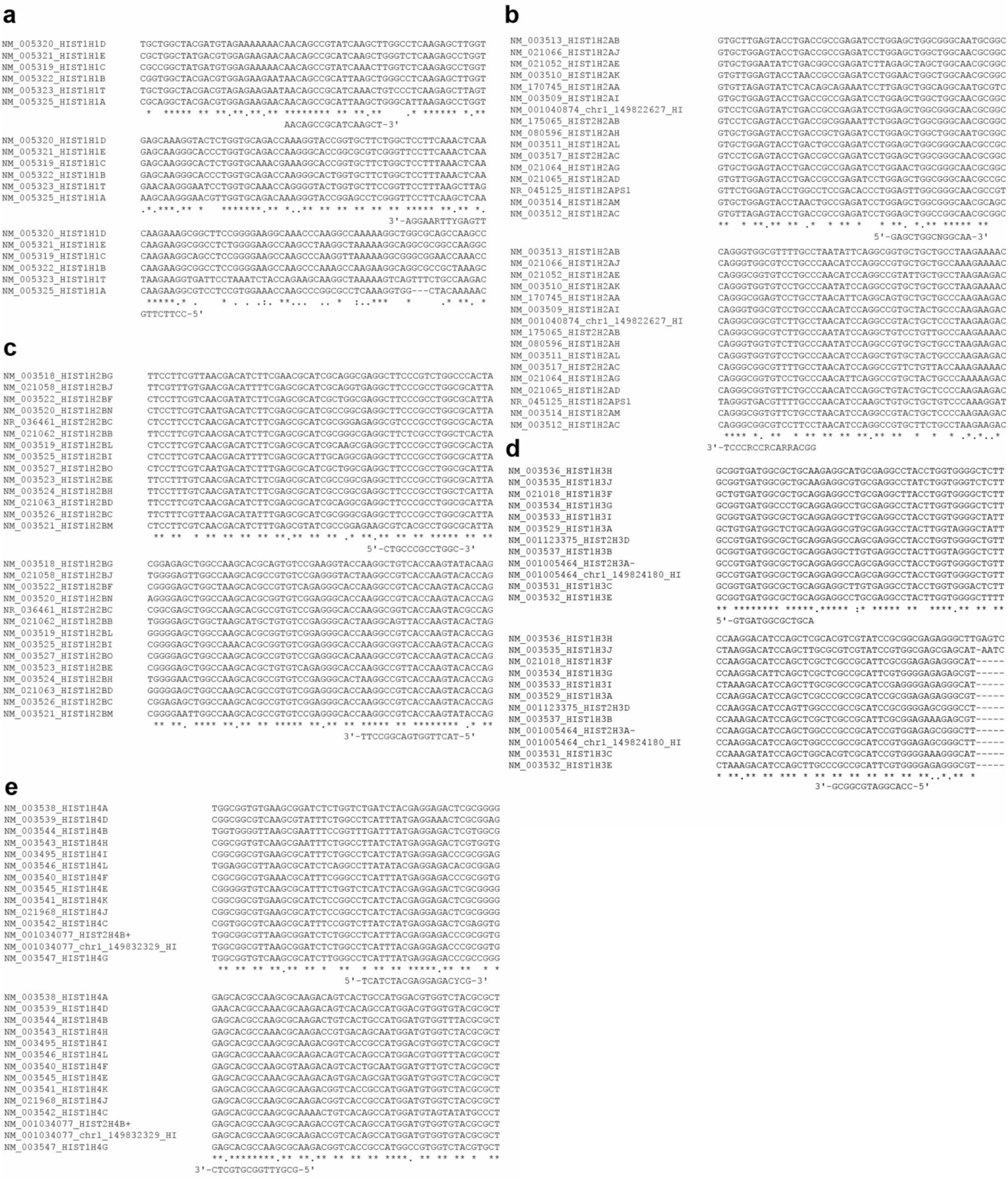
Replication-coupled histone gene alignments and primers. **(A-E)** Amplification primers targeting the most conserved regions of a Clustal-W alignment of histones H1, H2A, H2B, H3 and H4. Invariant (*) and conserved (.) residues are indicated below the alignment. Forward and reverse primers are displayed complementary to their coding sequence.

**Supplemental Figure 2.**
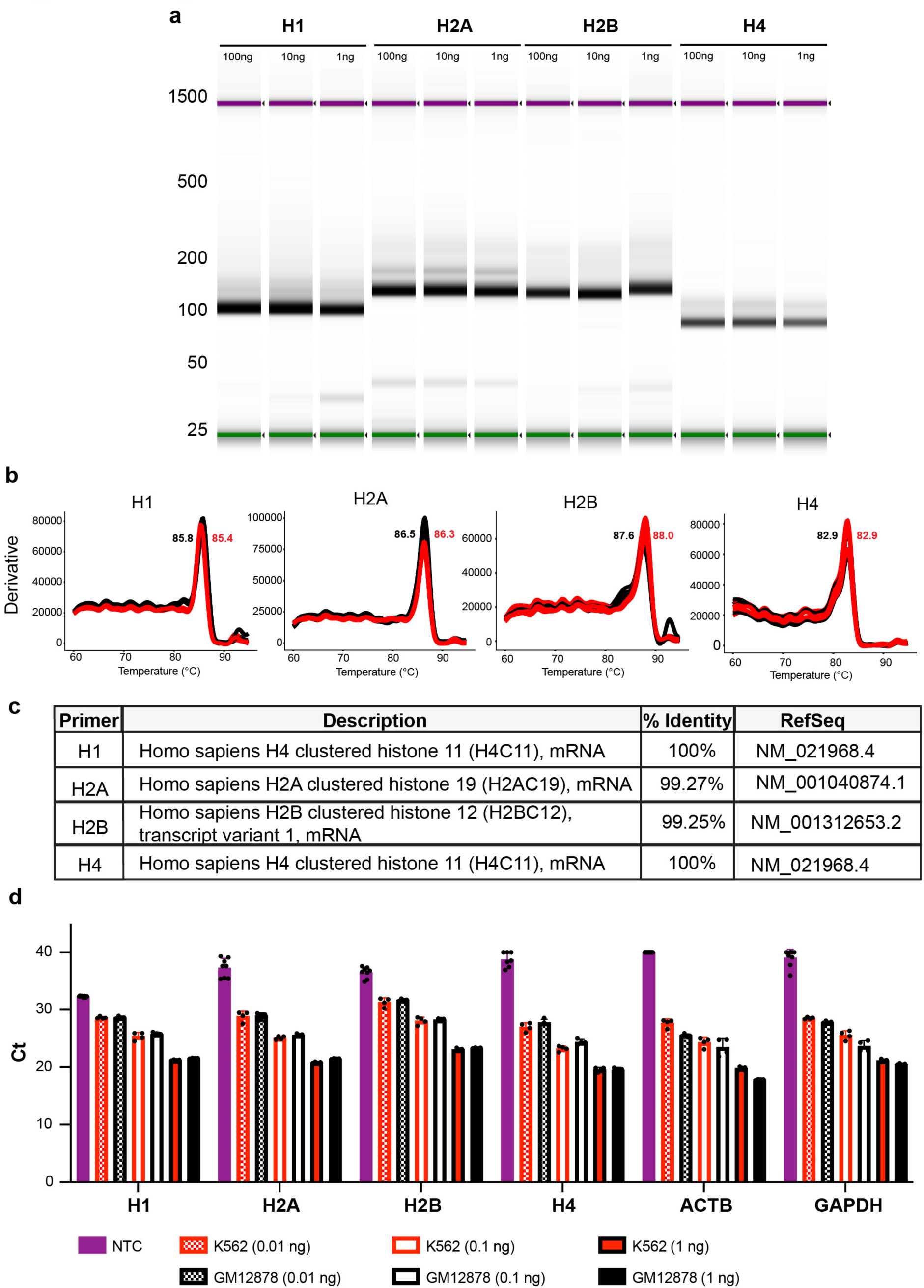
Validation of degenerate histone gene primers. **(A).** Automated capillary electrophoresis of PCR-amplified cDNA generated using degenerate histone gene primers. Lanes are grouped by target with each group representing a 100 ng, 10 ng, and 1 ng cDNA input series. **(B)** Derivative melt curves are shown for each primer set for 1 ng input cDNA (red = K562, black = GM12878). The melting temperature for each cell line was determined as the temperature at the maximum derivative signal and is annotated in each panel. **(C)** Consensus sequences were generated from amplification-free, single-molecule long-read sequencing of each amplicon. Table showing top BLASTn hit for each consensus read (NCBI core_nt, Homo sapiens). **(D)** Ct values for K562 and GM12878 cDNA (0.01 ng – 1ng) are compared to no template controls (NTC), where each point represents an individual technical replicate (n = 4-8), bars indicate mean with SD, and undetectable samples are assigned a Ct of 40.

**Supplemental Figure 3.**
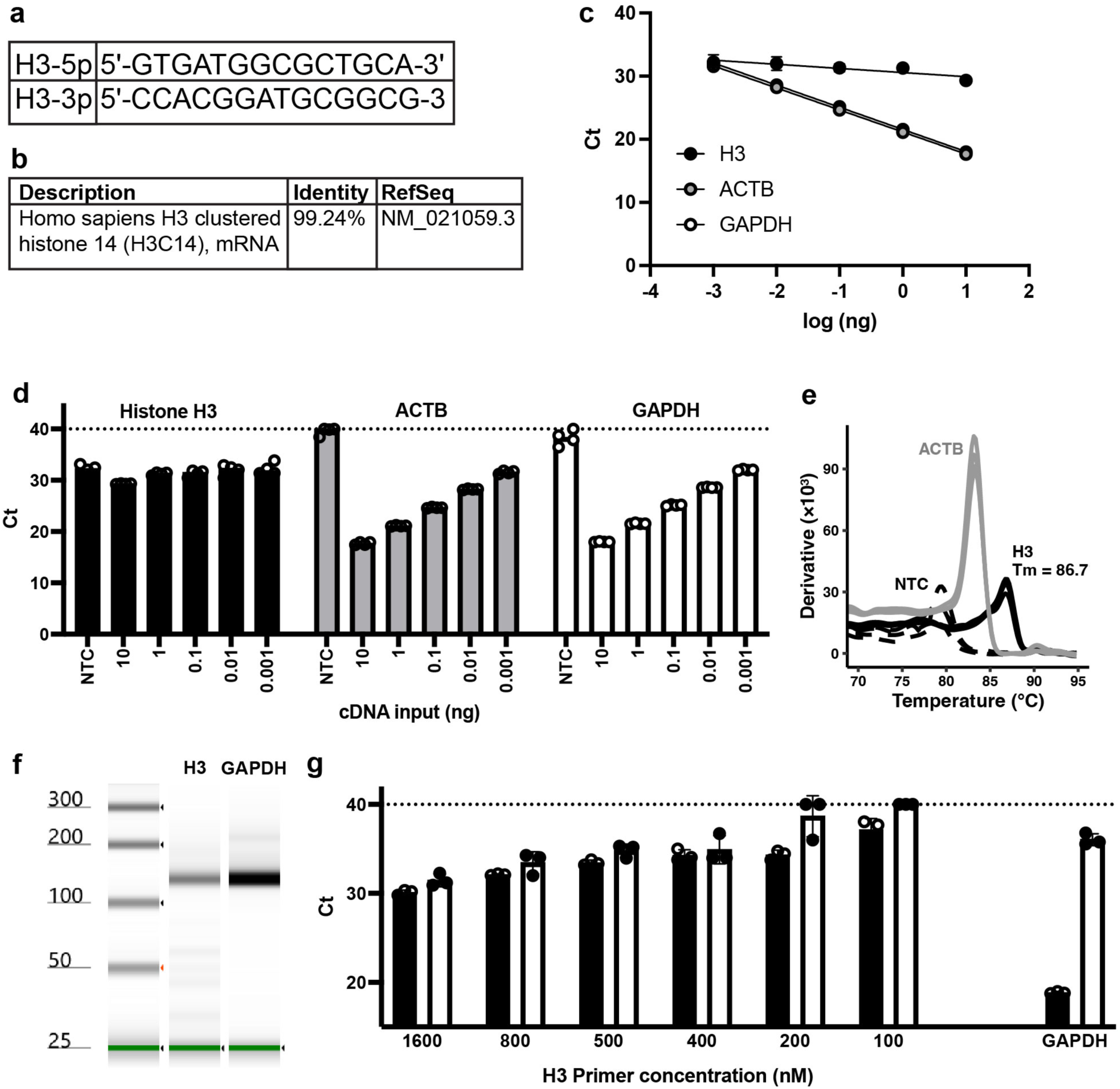
Characterization of histone H3 Primers. **(A).** Degenerate primers targeting the most conserved regions of histone H3. **(B)** Long-read sequencing of amplicons generated from K562 RNA by random hexamer priming and PCR using degenerate histone H3 primers identified H3_1 as the top BLASTn hit (NCBI core_nt, Homo sapiens). **(C,D)** qPCR was performed using serial dilutions (0.001–10 ng) of cDNA prepared by random hexamer-primed reverse transcription of K562 RNA. (C) Standard curves for histone H3 (black), ACTB (gray), and GAPDH (white) were generated from these reactions. Data points represent the mean of four technical replicates; error bars indicate SD, and solid lines represent linear regression fits for each primer set. (D) Ct values from reactions in (C) were plotted alongside no-template controls (NTCs). Each point represents an individual technical replicate (n = 4), and bars indicate the mean ± SD (undetectable reactions were assigned Ct of 40 shown by dotted line). **(E)** Derivative melt curve of reactions from (C). Traces are shown for histone H3 at 10 ng (solid black), no-template control (dashed), and ACTB (grey). **(F)** Automated capillary electrophoresis of PCR-amplified products from histone H3 and GAPDH primers from 10ng reactions in (C). **(G)** Ct values across histone H3 primer titration. qPCR performed on 5ng of random hexamer primed cDNA (template, black) using GAPDH and histone H3 primers (100 to 1600nM) compared to no-template controls (NTC, white). Bars indicate the mean of three technical replicates, error bars indicate SD, and individual replicates are plotted as points. No primer concentration suppressed background amplification with reliable detection of histone H3 product.

**Supplemental Figure 4.**
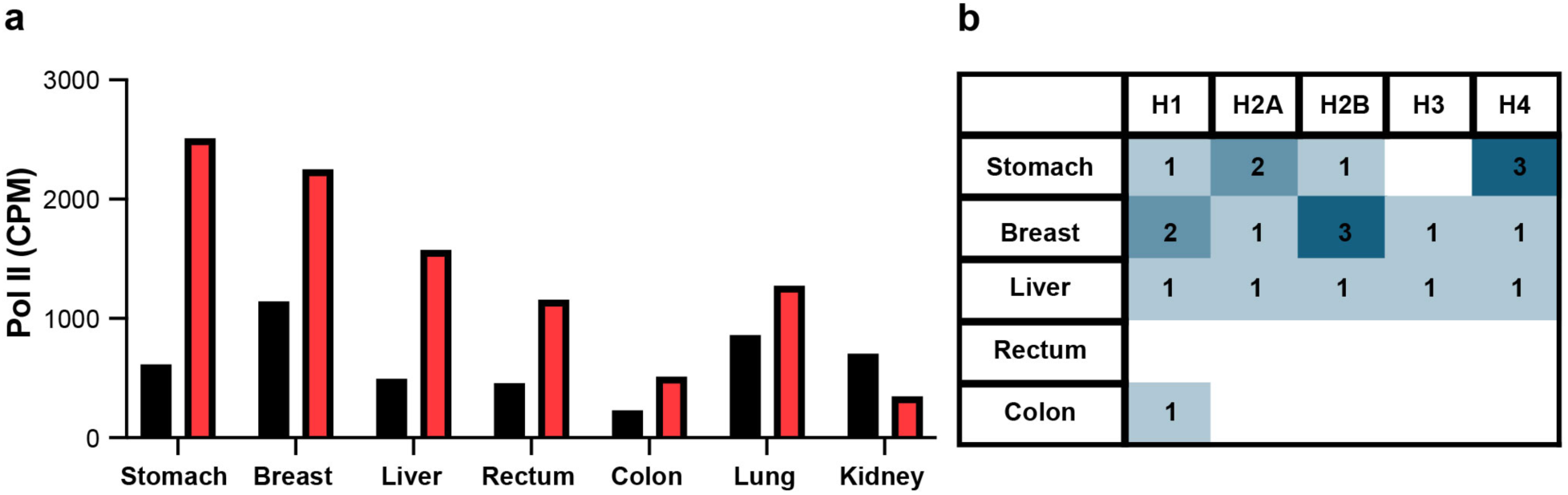
Individual histone genes are relatively decreased in tumors exhibiting cancer-associated hypertranscription and increased RNAPII occupancy at histone locus clusters. **(A)** RNAPII Signal at histone genes calculated as in (Fig. 3B) across multiple tumors and matched normal samples (black = normal, red = tumor) from our previous study (1). **(B)** Presence–absence matrix summarizing histone gene downregulation in samples that demonstrated genome-wide RNAPII hypertranscription. Each cell shows the number of histone genes decreased in the tumor sample relative to its matched normal.

**Supplemental Figure 5:**
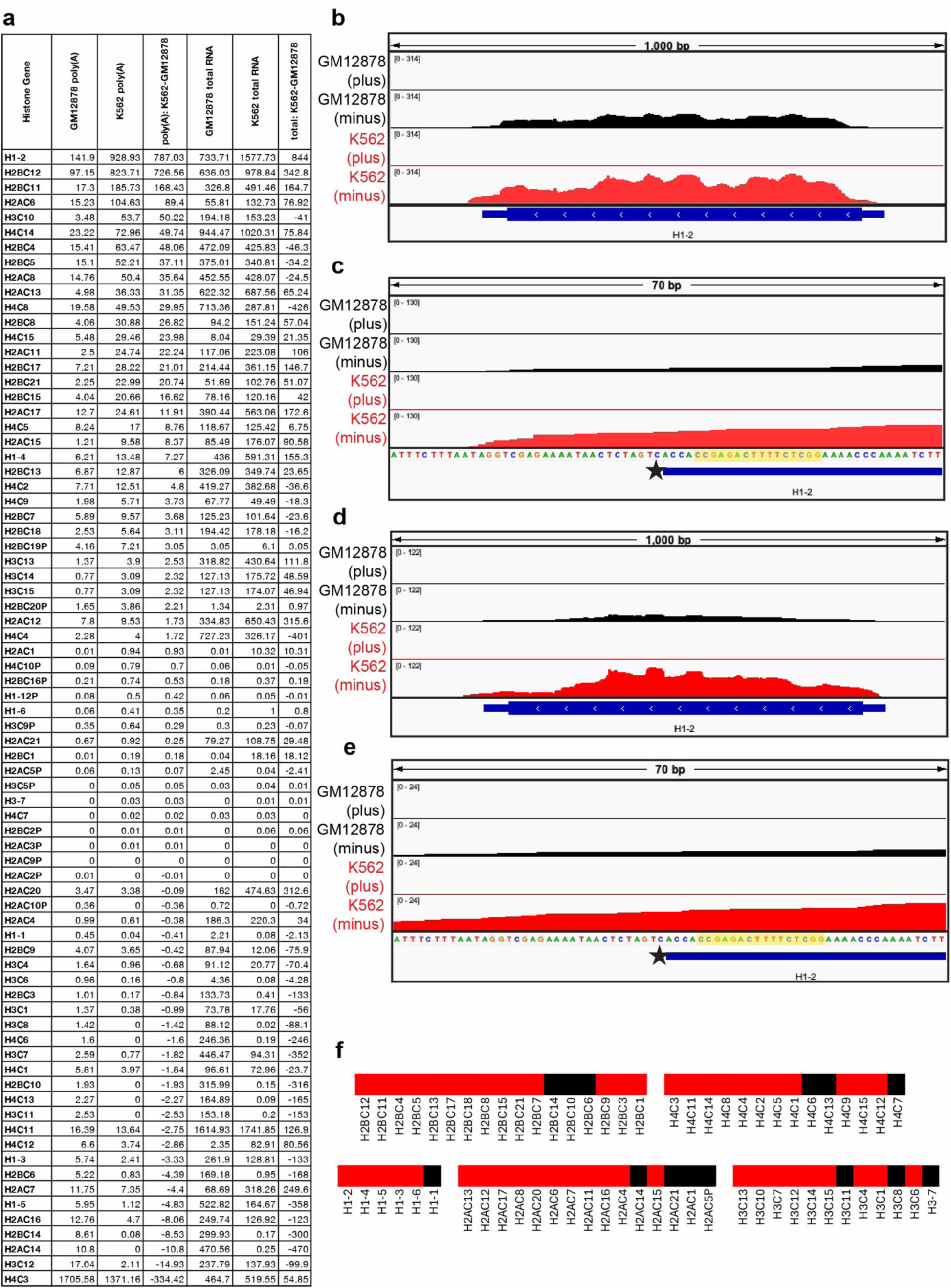
Evidence for polyadenylated histone mRNA in ENCODE RNA-seq. **(A)** Table of histone gene expression in GM12878 and K562 cells by RNA-seq library type. Mean transcripts per million (TPM) for histone genes from ENCODE gene-level quantifications of poly(A)+ RNA-seq (”GM12878 poly(A)”, “K562 poly(A)”) and rRNA-depleted total RNA-seq (”GM12878 total RNA”, “K562 total RNA”) libraries (accessions in Supplementary Table 1). TPM was averaged across replicates within each experiment, and experiment means were then averaged within each cell line and library type. The difference in mean TPM between K562 and GM12878 for each library type were calculated (”poly(A): K562-GM12878” and “total: K562-GM12878”). Genes are ordered by largest to smallest difference in poly(A) libraries. **(B, C)** Strand-specific signal of uniquely mapped reads from rRNA-depleted total RNA-seq data at H1-2, the top-ranked gene in (A), for GM12878 (black) and K562 (red). (B) Coverage across the gene. (C) Coverage at the 3ʹ end, with the conserved stem-loop highlighted in yellow, and the canonical cleavage site located 5 nt downstream of the stem-loop marked by a star. (ENCODE accessions: ENCFF509KPQ, ENCFF045ITU, ENCFF312ZLI, ENCFF530FJG). **(D, E)** Same as (B, C) for poly(A)+-selected libraries. (ENCODE accessions: ENCFF243ECE, ENCFF763UJG, ENCFF536ZVM, ENCFF839ZOT). In both library types, signal extends beyond the canonical cleavage site on the transcribed strand, consistent with a polyadenylated H1-2 transcript. **(F)** Heatmap of histone genes for presence of a polyadenylated isoform in K562 cells. Genes were scored positive (red) if (1) signal over the coding sequence was present in both poly(A)+ and total RNA-seq, and (2) total RNA-seq signal extended downstream of the canonical cleavage site on the transcribed strand.

**Supplemental Figure 6.**
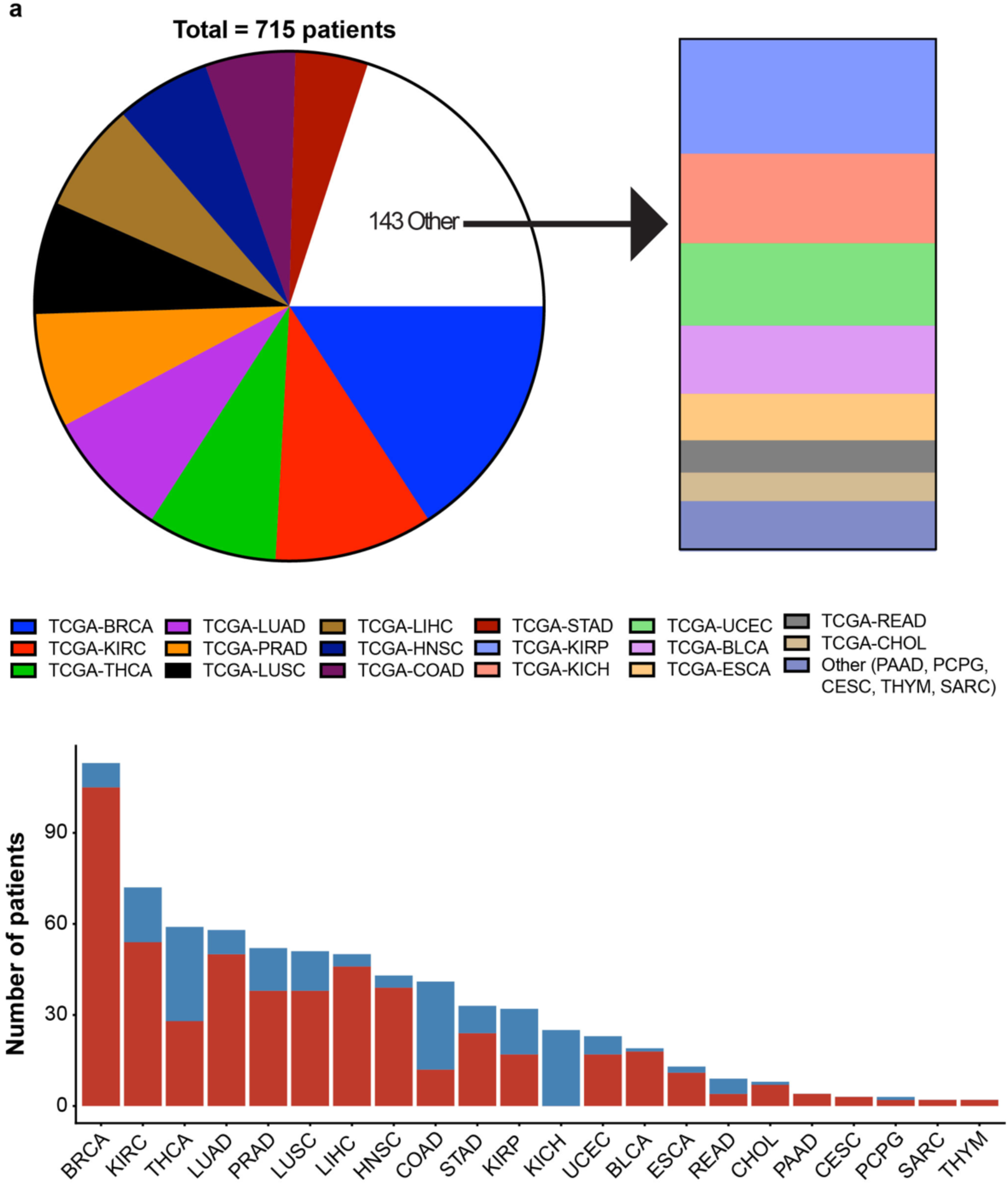
Distribution of Matched TCGA Patient Samples. **(A)** Composition of the study population across 22 TCGA projects (n=715 total patients). Pie chart highlights the 10 most frequent project cohorts: TCGA-BRCA (n=113), TCGA-KIRC (n=72), TCGA-THCA (n=59), TCGA-LUAD (n=58), TCGA-PRAD (n=52), TCGA-LUSC (n=51), TCGA-LIHC (n=50), TCGA-HNSC (n=43), TCGA-COAD (n=41), and TCGA-STAD (n=33). Stacked bar chart describes 12 additional projects: TCGA-KIRP (n=32), TCGA-KICH (n=25), TCGA-UCEC (n=23), TCGA-BLCA (n=19), TCGA-ESCA (n=13), TCGA-READ (n=9), TCGA-CHOL (n=8), TCGA-PAAD (n=4), TCGA-PCPG (n=3), TCGA-CESC (n=3), TCGA-THYM (n=2), and TCGA-SARC (n=2). **(B)** Polyadenylated histone gene expression is upregulated in the majority of tumor samples across nearly all cancer types examined. Stacked bar plots show the number of patients with higher (red) or lower (blue) summed histone TPM in primary tumor relative to matched normal tissue, for each of 22 TCGA cancer types.

**Supplemental Figure 7:**
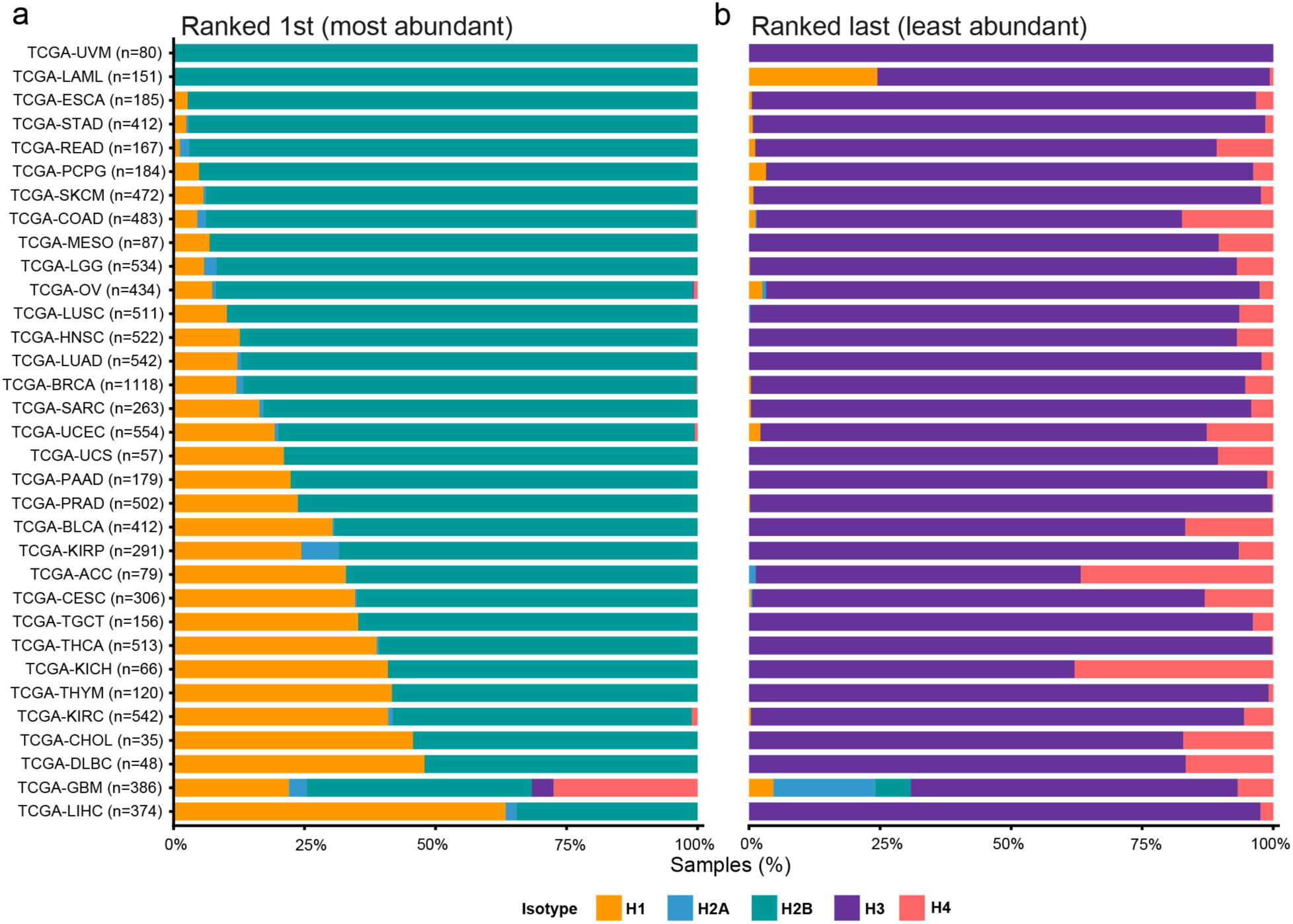
Pan-cancer histone isotype rank frequency across 33 TCGA cancer types. **(A, B)** Stacked horizontal bar charts showing the frequency of histone isotype ranking among 10,765 RNA-seq samples across 33 TCGA cancer types. For each sample, relative abundance of each histone isotype was calculated and then ranked from most abundant (highest, ranked first) to least abundant (ranked last). The proportion of samples in which each isotype ranked (A) first and (B) last were quantified for each cancer. Cancer types are ordered by decreasing H2B rank frequency.

**Supplemental Figure 8:**
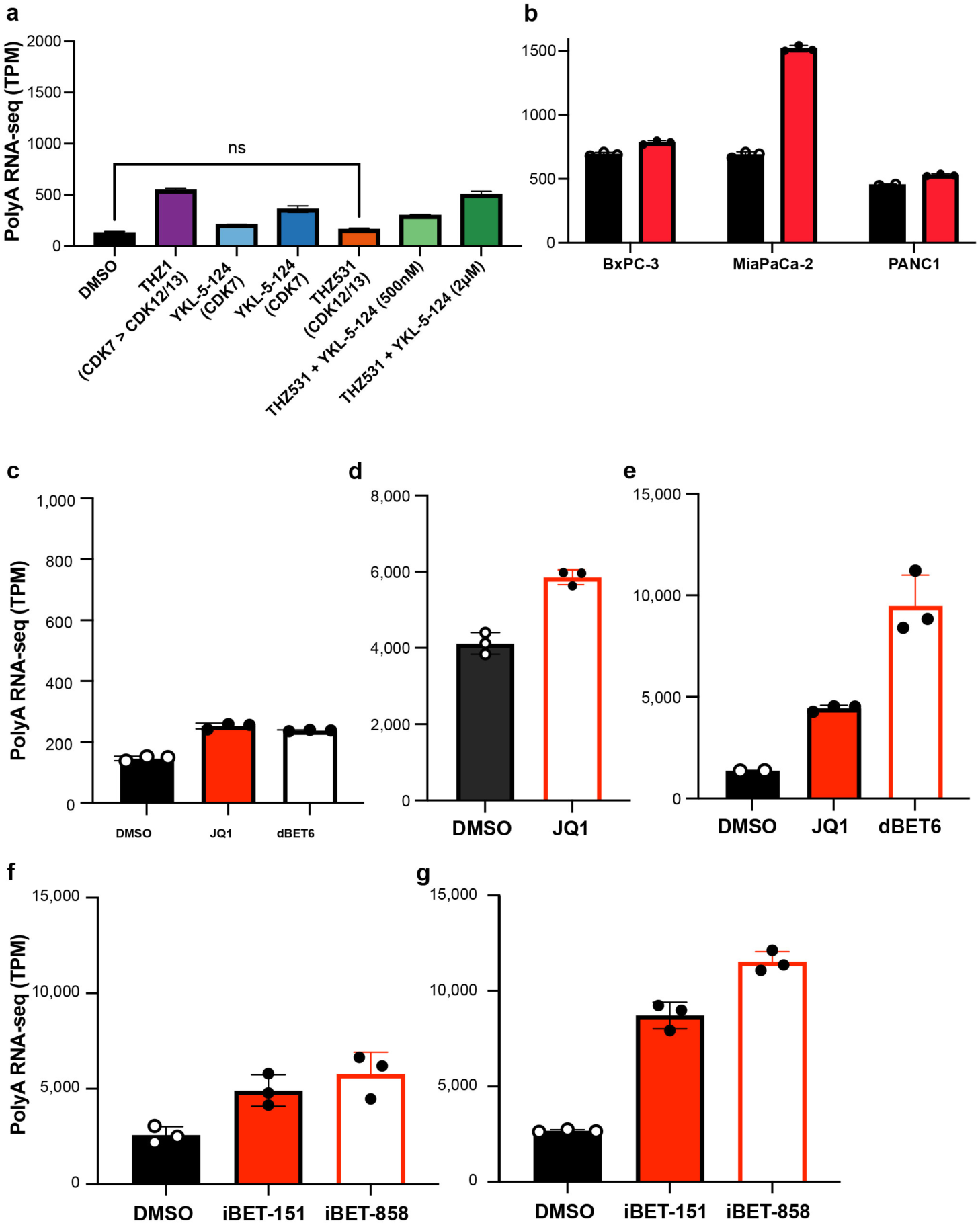
Chemical and genetic dissection of polyadenylated histone gene regulation. **(A)** HAP1 cells expressing mutant CDK7-C312S treated for 24 hours with DMSO, YKL-5-124 (CDK7-selective), THZ1 (CDK7/12/13), THZ531 (CDK12/13-selective), or THZ531 combined with YKL-5-124 (21). One-way ANOVA with Dunnett’s test; all comparisons significant (P adj ≤ 0.0007) except as noted (ns = not significant). **(B)** BxPC3, MiaPaCa-2 and PANC1 cells were treated with DMSO or THZ1 for 6hr (22). Unpaired t-test with Welch correction were run following Grubbs test comparing treated cells to control and were all found to be significantly greater in treatment (P ≤ 0.0031). **(C-E**) MOLT4 cells treated for 2h (C), T-ALL PDX for 6h (D), and primary T-ALL samples treated for 6h (E) with dBET6 or JQ1 (25). (C) One-way ANOVA with Dunnett’s test; all comparisons significant (P adj < 0.0001). (D) Unpaired two-tailed T-test (P = 0.0009). (E) one-way ANOVA with Dunnett’s test with all comparisons being statistically significant (P adj < 0.05). **(F-G)** OVCAR-3 ovarian cancer cells treated for (F) 4h and (G) 24h with either i-BET858 and i-BET151 (26). One-way ANOVA with Dunnett’s test; all comparisons statistically significant (P adj < 0.05).

## Reference

1. S. Henikoff et al., RNA polymerase II at histone genes predicts outcome in human cancer. Science 387, 737–743 (2025).

2. S. Cao et al., Estimation of tumor cell total mRNA expression in 15 cancer types predicts disease progression. Nat Biotechnol 40, 1624–1633 (2022).

3. M. Zatzman et al., Widespread hypertranscription in aggressive human cancers. Sci Adv 8, eabn0238 (2022).

4. Y. Zheng, K. Ahmad, S. Henikoff, Total whole-arm chromosome losses predict malignancy in human cancer. Proc Natl Acad Sci U S A 122, e2505385122 (2025).

5. W. F. Marzluff, K. P. Koreski, Birth and Death of Histone mRNAs. Trends Genet 33, 745–759 (2017).

6. D. B. Mahat et al., Single-cell nascent RNA sequencing unveils coordinated global transcription. Nature 631, 216–223 (2024).

7. K. Ahmad, S. Henikoff, Histone H3 variants specify modes of chromatin assembly. Proc Natl Acad Sci U S A 99 Suppl 4, 16477–16484 (2002).

8. W. F. Marzluff, P. Gongidi, K. R. Woods, J. Jin, L. J. Maltais, The human and mouse replication-dependent histone genes. Genomics 80, 487–498 (2002).

9. W. Albig, D. Doenecke, The human histone gene cluster at the D6S105 locus. Hum Genet 101, 284–294 (1997).

10. T. M. Rose et al., Consensus-degenerate hybrid oligonucleotide primers for amplification of distantly related sequences. Nucleic Acids Res 26, 1628–1635 (1998).

11. F. Traganos, L. Staiano-Coico, Z. Darzynkiewicz, M. R. Melamed, Effects of aclacinomycin on cell survival and cell cycle progression of cultured mammalian cells. Cancer Res 41, 2728–2737 (1981).

12. E. P. Consortium et al., Expanded encyclopaedias of DNA elements in the human and mouse genomes. Nature 583, 699–710 (2020).

13. M. P. Meers, D. Tenenbaum, S. Henikoff, Peak calling by Sparse Enrichment Analysis for CUT&RUN chromatin profiling. Epigenetics Chromatin 12, 42 (2019).

14. E. P. Consortium, An integrated encyclopedia of DNA elements in the human genome. Nature 489, 57–74 (2012).

15. M. S. Kagda et al., Data navigation on the ENCODE portal. Nat Commun 16, 9592 (2025).

16. Y. Luo et al., New developments on the Encyclopedia of DNA Elements (ENCODE) data portal. Nucleic Acids Res 48, D882–D889 (2020).

17. L. Zheng et al., Phosphorylation of stem-loop binding protein (SLBP) on two threonines triggers degradation of SLBP, the sole cell cycle-regulated factor required for regulation of histone mRNA processing, at the end of S phase. Mol Cell Biol 23, 1590–1601 (2003).

18. M. M. Koseoglu, L. M. Graves, W. F. Marzluff, Phosphorylation of threonine 61 by cyclin a/Cdk1 triggers degradation of stem-loop binding protein at the end of S phase. Mol Cell Biol 28, 4469–4479 (2008).

19. W. F. Marzluff, E. J. Wagner, R. J. Duronio, Metabolism and regulation of canonical histone mRNAs: life without a poly(A) tail. Nat Rev Genet 9, 843–854 (2008).

20. G. H. Cheng, A. Nandi, S. Clerk, A. I. Skoultchi, Different 3’-end processing produces two independently regulated mRNAs from a single H1 histone gene. Proc Natl Acad Sci U S A 86, 7002–7006 (1989).

21. G. H. Cheng, A. I. Skoultchi, Rapid induction of polyadenylated H1 histone mRNAs in mouse erythroleukemia cells is regulated by c-myc. Mol Cell Biol 9, 2332–2340 (1989).

22. S. M. Lyons et al., A subset of replication-dependent histone mRNAs are expressed as polyadenylated RNAs in terminally differentiated tissues. Nucleic Acids Res 44, 9190–9205 (2016).

23. G. F. Gao et al., Before and After: Comparison of Legacy and Harmonized TCGA Genomic Data Commons’ Data. Cell Syst 9, 24–34 e10 (2019).

24. R. J. Duronio, W. F. Marzluff, Coordinating cell cycle-regulated histone gene expression through assembly and function of the Histone Locus Body. RNA Biol 14, 726–738 (2017).

25. J. P. Kemp, Jr., et al., Cell-cycle-regulated transcriptional pausing of Drosophila replication-dependent histone genes. Mol Biol Cell 36, ar88 (2025).

26. W. Wu, J. E. Greene, K. Ahmad, S. Henikoff, KAS-CUT&Tag for direct mapping of transcription bubbles. bioRxiv 10.64898/2026.05.15.725569 (2026).

27. T. Velychko et al., CDK7 kinase activity promotes RNA polymerase II promoter escape by facilitating initiation factor release. Mol Cell 84, 2287–2303 e2210 (2024).

28. C. M. Olson et al., Development of a Selective CDK7 Covalent Inhibitor Reveals Predominant Cell-Cycle Phenotype. Cell Chem Biol 26, 792–803 e710 (2019).

29. P. Lu et al., THZ1 reveals CDK7-dependent transcriptional addictions in pancreatic cancer. Oncogene 38, 3932–3945 (2019).

30. L. Core, K. Adelman, Promoter-proximal pausing of RNA polymerase II: a nexus of gene regulation. Genes Dev 33, 960–982 (2019).

31. C. M. Olson et al., Pharmacological perturbation of CDK9 using selective CDK9 inhibition or degradation. Nat Chem Biol 14, 163–170 (2018).

32. G. E. Winter et al., BET Bromodomain Proteins Function as Master Transcription Elongation Factors Independent of CDK9 Recruitment. Mol Cell 67, 5–18 e19 (2017).

33. M. Quintela et al., Bromodomain inhibitor i-BET858 triggers a unique transcriptional response coupled to enhanced DNA damage, cell cycle arrest and apoptosis in high-grade ovarian carcinoma cells. Clin Epigenetics 15, 63 (2023).

34. H. Zheng et al., Identification of Integrator-PP2A complex (INTAC), an RNA polymerase II phosphatase. Science 370 (2020).

35. S. Hu et al., INTAC endonuclease and phosphatase modules differentially regulate transcription by RNA polymerase II. Mol Cell 83, 1588–1604 e1585 (2023).

36. A. Aakula et al., RAS and PP2A activities converge on epigenetic gene regulation. Life Sci Alliance 6 (2023).

37. F. Lu, et al., Integrator-mediated clustering of poised RNA polymerase II synchronizes histone transcription. bioRxiv 10.1101/2023.10.07.561364 (2024).

38. B. Stadelmayer et al., Integrator complex regulates NELF-mediated RNA polymerase II pause/release and processivity at coding genes. Nat Commun 5, 5531 (2014).

39. J. R. Skaar et al., The Integrator complex controls the termination of transcription at diverse classes of gene targets. Cell Res 25, 288–305 (2015).

40. K. Sabath et al., INTS10-INTS13-INTS14 form a functional module of Integrator that binds nucleic acids and the cleavage module. Nat Commun 11, 3422 (2020).

41. K. D. Sullivan, T. E. Mullen, W. F. Marzluff, E. J. Wagner, Knockdown of SLBP results in nuclear retention of histone mRNA. RNA 15, 459–472 (2009).

42. M. Krajewska et al., CDK12 loss in cancer cells affects DNA damage response genes through premature cleavage and polyadenylation. Nat Commun 10, 1757 (2019).

43. T. Narita et al., NELF interacts with CBC and participates in 3’ end processing of replication-dependent histone mRNAs. Mol Cell 26, 349–365 (2007).

44. J. J. Gruber et al., Ars2 promotes proper replication-dependent histone mRNA 3’ end formation. Mol Cell 45, 87–98 (2012).

45. J. Pirngruber et al., CDK9 directs H2B monoubiquitination and controls replication-dependent histone mRNA 3’-end processing. EMBO Rep 10, 894–900 (2009).

46. J. Pirngruber, A. Shchebet, S. A. Johnsen, Insights into the function of the human P-TEFb component CDK9 in the regulation of chromatin modifications and co-transcriptional mRNA processing. Cell Cycle 8, 3636–3642 (2009).

47. V. Kari et al., A subset of histone H2B genes produces polyadenylated mRNAs under a variety of cellular conditions. PLoS One 8, e63745 (2013).

48. S. He et al., SUMO2 promotes histone pre-mRNA processing by stabilizing histone locus body interactions and facilitating U7 snRNP assembly. Genes Dev 39, 1182–1197 (2025).

49. K. Ahmad, S. Henikoff, The histone variant H3.3 marks active chromatin by replication-independent nucleosome assembly. Mol Cell 9, 1191–1200 (2002).

50. C. Fletcher, N. Heintz, R. G. Roeder, Purification and characterization of OTF-1, a transcription factor regulating cell cycle expression of a human histone H2b gene. Cell 51, 773–781 (1987).

51. E. Griesbach, M. Schlackow, W. F. Marzluff, N. J. Proudfoot, Dual RNA 3’-end processing of H2A.X messenger RNA maintains DNA damage repair throughout the cell cycle. Nat Commun 12, 359 (2021).

52. C. Mannironi, W. M. Bonner, C. L. Hatch, H2A.X. a histone isoprotein with a conserved C-terminal sequence, is encoded by a novel mRNA with both DNA replication type and polyA 3’ processing signals. Nucleic Acids Res 17, 9113–9126 (1989).

53. P. B. Talbert, S. Henikoff, Histone variants at a glance. J Cell Sci 134 (2021).

54. D. Hanahan, Hallmarks of cancer-Then and now, and beyond. Cell 10.1016/j.cell.2025.12.049 (2026).

55. H. Weintraub, A possible role for histone in the synthesis of DNA. Nature 240, 449–453 (1972).

56. D. Chen et al., Cell cycle duration determines oncogenic transformation capacity. Nature 641, 1309–1318 (2025).

57. S. Giunta et al., CENP-A chromatin prevents replication stress at centromeres to avoid structural aneuploidy. Proceedings of the National Academy of Sciences of the United States of America 118 (2021).

58. M. D. Blower, B. A. Sullivan, G. H. Karpen, Conserved organization of centromeric chromatin in flies and humans. Dev Cell 2, 319–330 (2002).

59. J. A. Aguirre-Ghiso, J. J. Bravo-Cordero, W. Guo, G. Lauvau, M. S. Sosa, The sleeping threat: targeting cancer dormancy to transform metastasis therapy. Nat Rev Cancer 26, 513–533 (2026).

60. S. Henikoff et al., Epigenomic analysis of formalin-fixed paraffin-embedded samples by CUT&Tag. Nat Commun 14, 5930 (2023).

61. P. Channathodiyil, J. Houseley, Glyoxal fixation facilitates transcriptome analysis after antigen staining and cell sorting by flow cytometry. PLoS One 16, e0240769 (2021).

62. A. Colaprico et al., TCGAbiolinks: an R/Bioconductor package for integrative analysis of TCGA data. Nucleic Acids Res 44, e71 (2016).

63. L. G. Mascibroda et al., INTS13 variants causing a recessive developmental ciliopathy disrupt assembly of the Integrator complex. Nat Commun 13, 6054 (2022).

